# The CspC pseudoprotease regulates germination of *Clostridioides difficile* spores in response to multiple environmental signals

**DOI:** 10.1101/461657

**Authors:** Amy E. Rohlfing, Brian E. Eckenroth, Emily R. Forster, Yuzo Kevorkian, M. Lauren Donnelly, Hector Benito de la Puebla, Sylvie Doublié, Aimee Shen

## Abstract

The gastrointestinal pathogen, *Clostridioides difficile*, initiates infection when its metabolically dormant spore form germinates in the mammalian gut. While most spore-forming bacteria use transmembrane germinant receptors to sense nutrient germinants, *C. difficile* uses the soluble pseudoprotease, CspC, to detect bile salt germinants. To gain insight into CspC’s unique mechanism of action, we solved its crystal structure. Guided by this structure, we identified CspC mutations that confer either hypo- or hyper-sensitivity to bile salt germinant. Surprisingly, hyper-sensitive CspC variants exhibited bile salt-independent germination as well as increased sensitivity to amino acid and/or calcium co-germinants. Since the mechanism by which *C. difficile* spores sense co-germinants is unknown, our study provides the first evidence that CspC senses distinct classes of co-germinants in addition to bile salts. Since we observed that specific residues control CspC’s responsiveness to these different signals, CspC is critical for regulating *C. difficile* germination in response to multiple environmental signals.

## Introduction

*Clostridioides difficile*, more commonly known as *Clostridium difficile*, is a Gram-positive, spore-forming, obligate anaerobe that is the leading cause of health-care associated infections and gastroenteritis-associated death worldwide [1]. In the United States (US) alone, *C. difficile* causes over 500,000 infections per year, leading to ~29,000 deaths and over $5 billion in medical costs [2]. While immunocompetent individuals are usually protected from *C. difficile* infection by the intestinal microflora, antibiotic treatment can render individuals susceptible to *C. difficile* infections due to disruption of the protective gut microbiota [3-5].

*C. difficile* infections are characterized by high rates of disease recurrence: approximately one in five patients that recover from a *C. difficile* infection will acquire a second infection within three months [2, 6, 7]. Both *C. difficile*’s vegetative cell and spore form contribute to recurrent infections [1, 8]: the vegetative form of *C. difficile* antagonizes growth of the protective microbiota by producing inflammation-inducing toxins [5], while its dormant spore form, which can resist commonly used disinfectants and harsh conditions [9, 10], allows *C. difficile* to outlast antibiotic treatment and persist in the environment for long periods of time [11].

Since the vegetative form of *C. difficile* cannot survive outside the anaerobic environment of the gastrointestinal tract, *C. difficile*’s aerotolerant, dormant spore form is its major infectious particle [9]. Consequently, *C. difficile* infections depend upon *C. difficile* spore germination. When ingested spores reach the small intestine, they sense mammalian-specific bile salts, which initiate a signaling cascade that allows spores to exit dormancy during germination [12-14]. Germinating spores outgrow to form the vegetative, toxin-producing cells that are responsible for *C. difficile* disease symptoms, which can range from severe diarrhea to pseudomembraneous colitis, toxic megacolon, and death [1, 6]. Since germination is required for *C. difficile* to initiate infection, therapeutics that inhibit germination to prevent *C. difficile* infection are currently being developed [15-18]. However, the molecular mechanisms underlying the *C. difficile* germination signaling cascade are poorly understood.

Indeed, recent studies indicate that *C. difficile*’s germination pathway is unique among other spore forming organisms [19, 20]. Almost all spore-forming organisms studied to date encode transmembrane germinant receptors of the Ger family [21, 22], which sense nutrient germinants, like amino acids, nucleic acids, and sugars [23]. In contrast, *C. difficile* does not encode Ger family receptors. Furthermore, the primary germinants for *C. difficile* are cholate-derived bile salts (especially taurocholate [24]), which are not known to be a nutrient source. Instead, genetic data suggest that *C. difficile* uses a soluble pseudoprotease, CspC, to directly sense bile salts [12]. CspC was identified in a genetic screen for mutants that germinate in response to the bile salt, chenodeoxycholate, which typically acts as a competitive inhibitor of germination [12, 25]. Remarkably, a single point mutation in CspC (glycine 457 to arginine, G457R) expanded *C. difficile*’s germinant specificity to permit germination in response to chenodeoxycholate as well as taurocholate. Since point mutations in CspC that prevent spore germination were also identified, CspC has been proposed to be the bile salt germinant receptor [12].

Interestingly, *C. difficile* CspC is a catalytically inactive member of the Csp family of proteases, which were first identified in *Clostridium perfringens* as being responsible for proteolytically activating the cortex hydrolase, SleC [26, 27]. Active SleC degrades the thick protective cortex layer [28, 29], a step that is essential for all spores to exit dormancy [23]. *C. perfringens* strains encode one or more of three Csps, CspA, CspB, and CspC [30, 31], which apparently have redundant functions during germination [31]. We previously solved the crystal structure of CspB from *C. perfringens* and confirmed that Csps are structurally similar to other subtilisin-like serine proteases [32, 33]. CspB protease activity depends on a catalytic triad consisting of Asp, His, and Ser, and the prodomain of CspB acts as an intramolecular chaperone that is auto-processed upon proper folding of CspB’s subtilisin-like serine protease domain. However, unlike other subtilisin-like serine proteases studied to date, the prodomain of CspB stays associated with the subtilase domain after cleavage and sterically occludes CspB’s active site [32].

While C*. difficile* strains encode homologs of all three Csp proteins, only CspB has an intact Asp, His, and Ser catalytic triad. CspA and CspC carry mutations in two of the catalytic residues, rendering them pseudoproteases [30]. As a result, only CspB can proteolytically activate SleC during *C. difficile* spore germination [32]. Furthermore, CspB is produced as a fusion to CspA in sporulating *C. difficile* cells. The CspBA fusion protein undergoes interdomain processing such that CspB and CspA are present as separate domains in mature spores [30, 32]. Notably, the CspBA fusion protein and the pseudoprotease nature of CspA and CspC appear to be unique to *C. difficile* and the Peptostreptococcaceae family [34]. In contrast, members of the Clostridiaceae and Lachnospiraceae family exclusively produce Csp family proteases as individual proteins with intact catalytic triads [30].

Both the CspA and CspC pseudoproteases play unique roles in regulating *C. difficile* spore germination. CspA controls CspC levels in spores, controlling CspC’s incorporation and/or stability in mature spores, and as described earlier, CspC is the proposed germinant receptor. Further underscoring the uniqueness of *C. difficile*’s germination pathway, we recently identified a *C. difficile*-specific protein, GerG, as a regulator of Csp family members’ incorporation into spores [35].

To gain insight into the molecular mechanisms by which *C. difficile* spores sense germinant, we solved the crystal structure of *C. difficile* CspC to 1.55 Å resolution. This structure revealed unique features of the *C. difficile* CspC pseudoprotease when compared to the *C. perfringens* CspB protease. Structure-function analyses identified several flexible residues critical for CspC function that confer either hyper- or hypo-sensitivity to bile salt germinant. Further analyses revealed that some of these mutations alter sensitivity to amino acid and/or calcium co-germinants, which potentiate bile salt-induced germination in *C. difficile*. Since the mechanism by which co-germinants are sensed by *C. difficile* spores was previously unknown [19, 20], our study reveals for the first time that *C. difficile* CspC integrates multiple germinant and co-germinant signals to induce spore germination and raises important new questions regarding how these signals are specifically sensed and transduced.

## Results

### Overall structure of CspC

To determine how *C. difficile* CspC directly senses bile salt germinants, we attempted to crystallize recombinant C-terminally His_6_-tagged CspC (wild-type or the G457R variant) alone or in the presence of either taurocholate or chenodeoxycholate. While crystals were observed in all three conditions using wild-type CspC, no crystals were obtained in our screening with the G457R variant. Furthermore, diffraction quality crystals were only obtained with wild-type CspC in the absence of bile salts. Determination of the crystal structure of the *C. difficile* CspC pseudoprotease revealed that it shares a conserved three domain architecture with the *C. perfringens* CspB protease (**Fig. 1A**), consisting of an N-terminal prodomain, subtilase domain, and jelly roll domain (**Fig. 2A**, [32]). The jelly roll is a ~130 amino acid insertion in the subtilase domain that forms a β-barrel domain that appears unique to the Csp family of subtilisin like-serine proteases. We previously determined that the jelly roll domain provides CspB with remarkable structural rigidity, conferring thermostability and resistance to proteolysis [32]. In both CspB and CspC, the jelly roll domain emerges from the subtilase domain and extensively interacts with both the subtilase domain and the prodomain. Overall, the secondary structure of these proteins is highly similar (**Fig. 1B**), with a 1.03 Å rmsd over 1200 atoms of the backbone trace.

**Figure 1.**
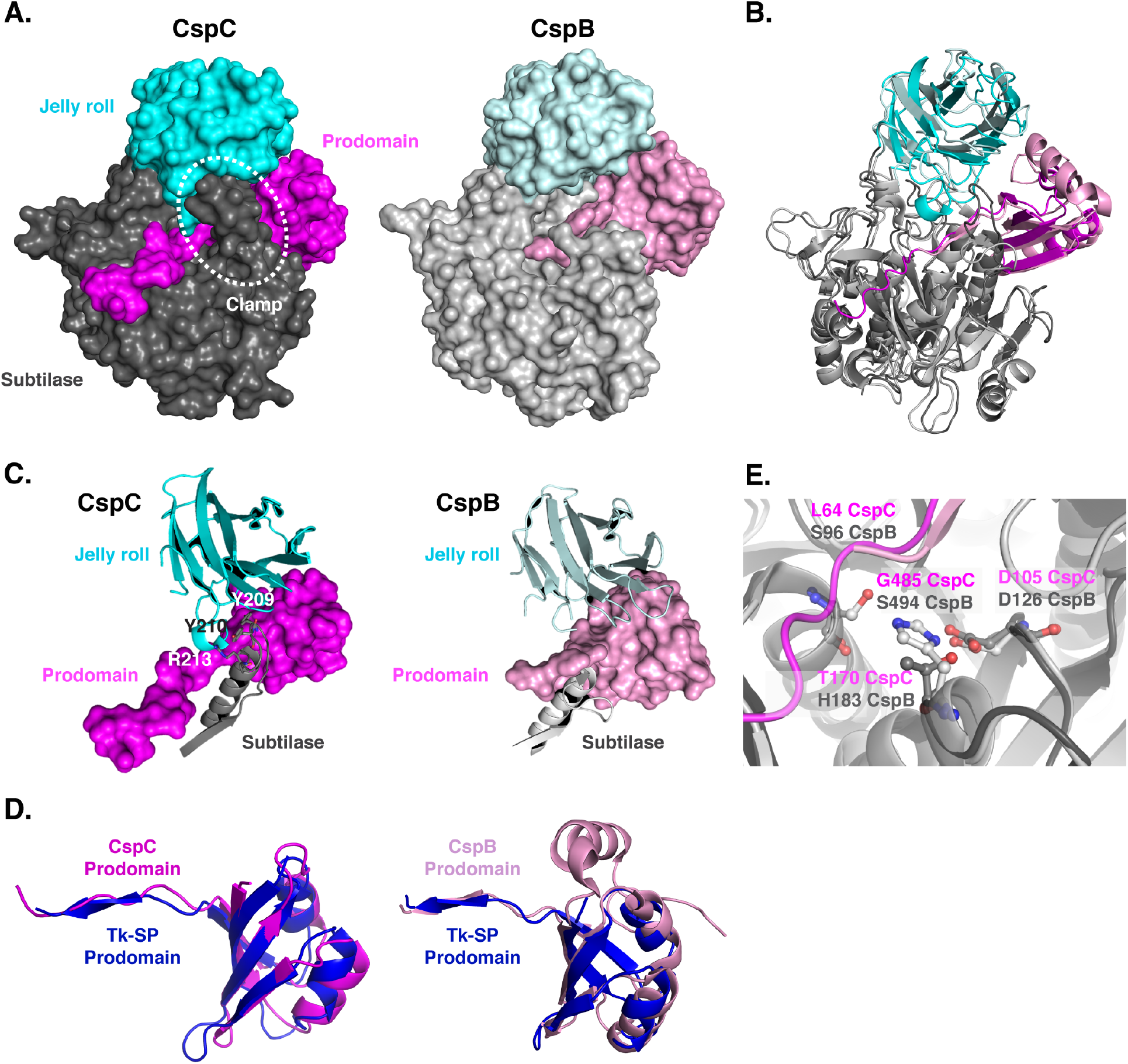
Structural comparison of *C. difficile* CspC to *C. perfringens* CspB. (A) Space-fill models of CspC and CspB [32] with the respective prodomains colored in pink, subtilase domains in grey, and jell roll domains in cyan (lighter tones denote CspB). The prodomain “clamp” in CspC formed by interactions between the jelly roll domain and subtilase domains is highlighted. (B) Ribbon representation overlay of CspC and CspB structures. A least squares superposition of the subtilase domain reveals a 1.03 Å rms over 1200 atoms of the backbone trace. (C) Comparison of the CspC prodomain “clamp” relative to CspB. The jelly roll domain interacts with a clamping loop from the CspC subtilase domain through which the CspC prodomain is threaded. Specific residues involved in the CspC jelly roll-subtilase domain interactions are marked. This interaction is absent in CspB due to a shorter subtilase domain loop, which is in a different orientation, and the disorder of the jelly roll domain interacting loop. (D) Superposition of the CspC and CspB prodomains, respectively, with the canonical prodomain from Tk-subtilisin [36] (blue). (E) Overlay of the CspB catalytic triad (S494, H183 and D126) to CspC (G485, T170 and D105). The position of the terminal residue of the CspB prodomain (S96) as well as the equivalent residue in CspC’s uncleaved prodomain (L64) is indicated.

Despite this shared secondary structure, *C. difficile* CspC does not autoprocess its prodomain due to substitutions in its catalytic triad. In addition, CspC’s prodomain is more closely associated with the subtilase and jelly roll domains due to a clamping loop shown in the surface representation (**Fig. 1A**). This loop resides between beta strand 4 and helix 4 within the subtilase domain (residues 203-216) and interacts with the jelly roll domain via packing of tyrosine 209 and tyrosine 210 as well as hydrogen bonds from arginine 213 to backbone carbonyls in the 402-407 residue loop of the jelly roll domain (**Fig. 1C**). In CspB, the equivalent subtilase domain loop is shorter (residues 214-225) and in a different conformation, and the jelly roll domain loop is disordered, possibly due to the autoprocessing of the prodomain by CspB’s subtilase domain (**Fig. 1C**).

The prodomain of CspC aligns more closely with the canonical Tk-SP subtilisin prodomain [36] because CspB’s prodomain is larger due to a helical insert (**Fig. 1D**).

Nevertheless, both prodomains share a similar orientation relative to active (or degenerate) site residues (**Fig. 1E**). As a result, leucine 64 of CspC (**Fig. 1E**) is equivalent to the autocleavage site of the CspB *perfringens* prodomain, serine 96 [27, 32]), and would be positioned to undergo autoprocessing if CspC’s catalytic triad were intact. Notably, the degenerate site residues of CspC’s pseudotriad share the same orientation as the catalytic triad of CspB and other subtilisin-like serine proteases (**Fig. 1E**).

### Restoring the catalytic triad to CspC disrupts protein folding and function

This latter observation prompted us to test whether CspC could be converted into an active protease that could undergo autoprocessing like other subtilisin-like serine proteases [33]. To this end, we restored CspC’s catalytic triad by cloning complementation constructs encoding amino acid substitutions of threonine 170 to histidine (T170H) and glycine 485 to serine (G485S) both individually and in combination. These constructs, along with all other *cspC* constructs analyzed in this manuscript, were expressed from the native *cspBA-cspC* promoter and integrated into the *pyrE* locus of Δ*cspC*Δ*pyrE* using allele-coupled exchange [37]. To assess whether the T170H-G485S substitutions activated CspC autoprocessing activity, we analyzed CspC in sporulating cells using western blotting. Notably, no CspC autoprocessing was detected with the T170H/G485S (2xcat) double mutant; in fact, no CspC was detectable in sporulating cell lysates of this mutant strain (**Fig. 2B**). Furthermore, CspC levels were markedly decreased in the G485S mutant and slightly reduced in the T170H mutant, implying that these mutations reduce CspC production and/or stability in sporulating cells. No decrease in CspBA levels was observed in any of the mutant strains (**Fig. 2B**), consistent with the observation that CspC does not affect CspBA [38].

**Figure 2.**
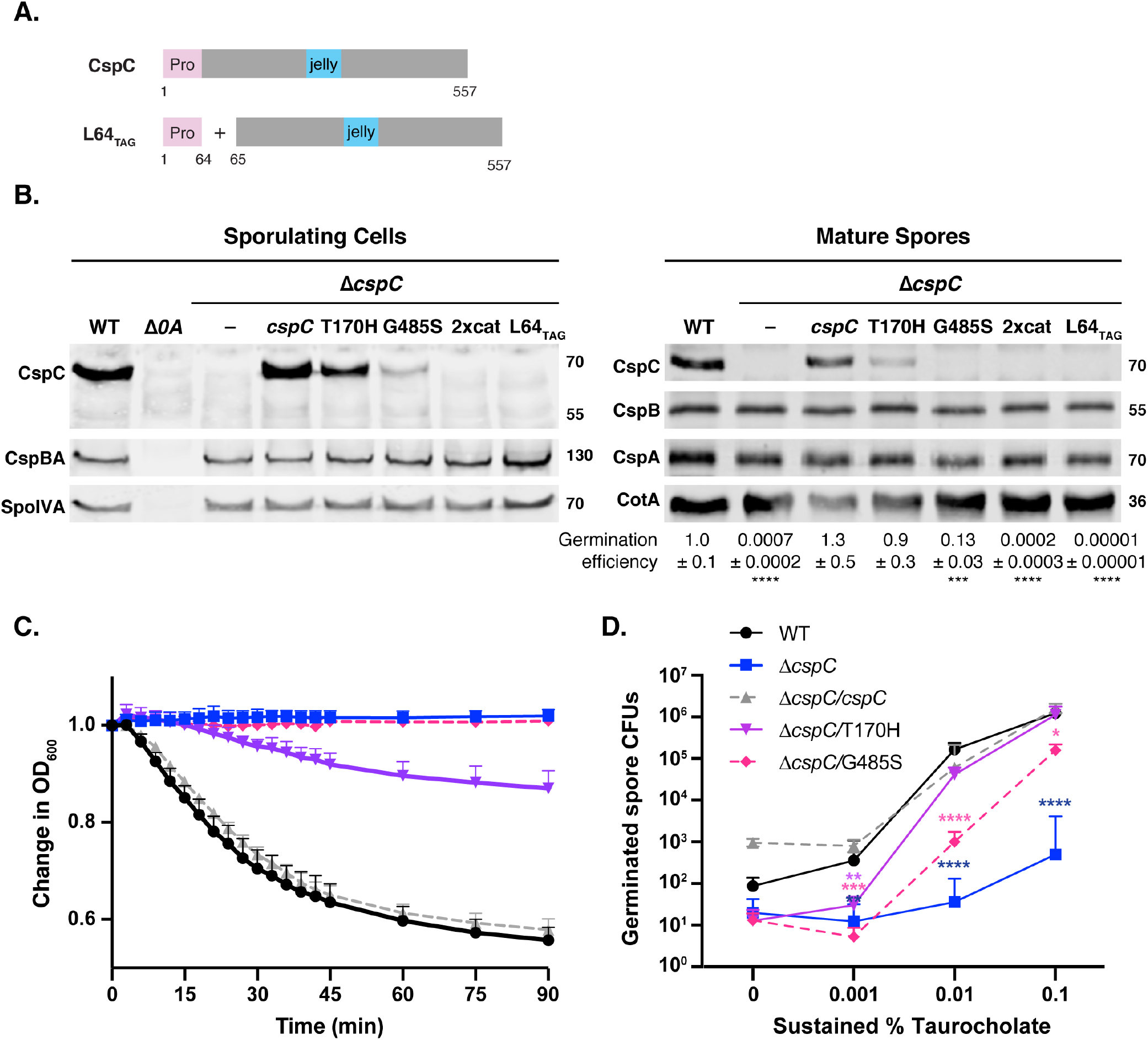
Mutations to restore CspC’s catalytic triad decrease CspC levels and germination efficiency. (A) Schematic of wild-type CspC and a construct encoding the prodomain *in trans* (L64_TAG_). “Pro” denotes the prodomain; “jelly” indicates the jelly roll domain; and the grey rectangle represents the subtilase domain. L64_TAG_ encodes a variant in which the CspC prodomain is produced *in trans* from the remainder of CspC through the introduction of a stop codon after codon 64 and a ribosome binding site and start codon before codon 65. (B) Western blot analyses of CspC and CspB(A) in sporulating cells and purified spores from wild type 630Δ*erm*-p, Δ*cspC*, and Δ*cspC* complemented with either wild-type *cspC* or *cspC* variants. 2xcat encodes a T170H-G485S double mutant. Δ*spo0A* (Δ*0A*) was used as a negative control for sporulating cells. SpoIVA was used as a loading control for sporulating cells, while CotA was used as a loading control for purified spores. Full-length CspBA detected by anti-CspB antibodies is shown for sporulating cells, while the individual CspB and CspA domains detected by anti-CspB and anti-CspA antibodies, respectively, are shown for purified spores. The germination efficiency of spores plated on BHIS media containing 0.1% taurocholate is also shown relative to wild type. The mean and standard deviations shown are based on three biological replicates performed on two independent spore purifications. Statistical significance relative to wild type was determined using a one-way ANOVA and Tukey’s test. **** p < 0.0001, *** p < 0.001. (C) Optical density (OD_600_) analyses of spore germination over time in CspC degenerate site single mutants. Purified spores from the indicated strains were incubated in BHIS with and without 1% taurocholate, and the OD_600_ of the samples was monitored using a spectrophotometer. The change in OD_600_ represents the ratio of the OD_600_ of the treated to the untreated sample relative to the ratio at time zero. The averages of the results from four biological replicates performed on at least two independent spore preps are shown. The error bars indicate the standard deviation for each timepoint measured. (D) Germinant sensitivity of degenerate site mutant spores plated on BHIS containing increasing concentrations of taurocholate. The number of colony forming units (CFUs) produced by germinating spores is shown. The mean and standard deviations shown are based on three biological replicates performed on two independent spore purifications. Statistical significance relative to wild type was determined using a one-way ANOVA and Tukey’s test. **** p < 0.0001, *** p < 0.001, ** p < 0.01, * p < 0.05.

To assess whether the reduced CspC levels in the CspC degenerate site mutants were due to general protein folding defects, we cloned these alleles into recombinant protein expression vectors and measured CspC-His_6_ production and purification levels in *E. coli*. Notably, recombinant CspC_T170H-G485S_ did not autoprocess when produced in *E. coli* (**Fig. S1B**), indicating that restoring the catalytic triad does not reconstitute CspC protease activity. Consistent with the hypothesis that mutations in the degenerate active site impair proper CspC folding in *C. difficile*, recombinant CspC_T170H_, CspC_G485S_, and CspC_T170H/G485S_ were purified from the soluble fraction at a much lower efficiency than wild-type CspC-His_6_ (Elution, **Fig. S1B**) even though all three strains exhibit wild-type levels upon IPTG induction (Induced, **Fig. S1B**). Unfortunately, since most of recombinant CspC produced in *E. coli* was insoluble, it was difficult to assess whether the different mutants produced higher levels of insoluble protein.

To evaluate whether the decreased CspC levels observed in the *C. difficile* degenerate site mutants impact spore germination, we measured the ability of these mutant alleles to complement Δ*cspC*’s germination defect. Purified spores from wild-type, Δ*cspC*, Δ*cspC*/*cspC*, and the degenerate site mutant complementation strains were plated on BHIS media containing 0.1% taurocholate germinant, and the number of colony forming units (CFUs) that arose from germinating spores relative to wild type was determined. The T170H allele restored germination to wild-type levels even though CspC_T170H_ protein levels were visibly decreased in western blot analyses of purified spores (**Fig. 2B**). In contrast, G485S mutant spores exhibited a ~10-fold defect in germination efficiency even though CspC_G485S_ protein was barely detectable. The T170H/G485S double mutant’s germination defect (~1,000-fold) was the most severe, resembling that of the parental Δ*cspC* strain, consistent with the absence of detectable CspC_T170H/G485S_ in sporulating cells and purified spores (**Fig. 2B**). Similar to the phenotypes of other germination-receptor mutants [35, 38-40], Δ*cspC* spores exhibit low levels of “spontaneous” germination.

Although the G485S mutant’s germination defect was only ~10-fold when spores were plated on BHIS containing 0.1% germinant, G485S colonies were slower to appear than colonies arising from wild-type spores. To test whether the G485S and T170H degenerate site mutations affected the rate of germination, we used an optical density assay to monitor early germination events. This assay measures the decrease in optical density of a population of germinating spores over time due to cortex hydrolysis and core hydration [41]. Whereas wild-type spores decrease in optical density by ~40%, the optical density of the G485S mutant did not appreciably change over the 3 hr assay period, similar to the Δ*cspC* strain (**Fig. 2C**). Delayed germination was observed in the T170H mutant relative to both wild type and the wild-type *cspC* complementation strains even though *cspC*_T170H_ did not exhibit a germination defect in the plate-based CFU assay. Taken together, these results indicate that single mutations in CspC’s degenerate active site impair folding and/or decrease CspC stability in sporulating cells and slow down spore germination rates. Restoring the full catalytic triad to CspC fails to activate CspC catalytic activity and significantly reduces CspC protein levels in sporulating cells.

Since the degenerate site mutants produced wild-type levels of both CspBA in sporulating cells and CspB and CspA in purified spores (**Fig. 2B**), the observed germination defects are due to impaired CspC function and/or decreased protein levels. We wondered whether the reduced CspC levels in the T170H and G485S variants would decrease germinant sensitivity, since decreased CspB, CspA, and CspC levels in *gerG* mutant spores diminish the responsiveness of *C. difficile* spores to germinant [35]. To measure germinant sensitivity, we plated *cspC* mutant spores on rich media with varying concentrations of taurocholate germinant. On plates containing 0.001% taurocholate, CspC_T170H_ and CspC_G485S_ mutant spores germinated to a similar extent as Δ*cspC* spores and at lower levels than wild-type spores (p ≤ 0.005, **Fig. 2D**). However, on plates with 0.01% taurocholate, T170H mutant spores germinated to near wild-type levels, whereas G485S spores exhibited an ~100-fold decrease relative to wild type. At the highest concentration of germinant tested (0.1% taurocholate), CspC_T170H_ mutant spores were indistinguishable from wild-type spores, and the CspC_G485S_ mutant spores had an ~1-log defect in CFUs consistent with our prior findings (**Fig. 2B**). Thus, decreased CspC protein levels and/or function in the degenerate site mutants impairs germinant sensing even when CspB and CspA are present at wild-type levels, consistent with the idea that CspC is the germinant receptor.

### Loss-of-function CspC mutations identified in a genetic screen cluster to the degenerate active site region

The CspC structure also allowed us to localize the CspC loss-of-function mutations identified by Francis *et al*. in their EMS mutagenesis screen for *C. difficile* germination mutants [12]. Of the six CspC mutations identified, three clones carried substitutions of glycine 171 to arginine (G171R), two clones had substitutions of glycine 483 to arginine (G483R), and individual clones carried substitutions of either serine 488 to asparagine (S488N) or glycine 276 to arginine (G276R). The final mutant carried two point mutations: valine 272 to glycine and serine 443 to asparagine (V272G/S443N). Notably, 5 out of 6 of these residues cluster to the degenerate active site region and are oriented towards the prodomain (**Fig. S1A**), while the other residue, serine 443, was mutated in combination with a residue close to the degenerate site, V272.

Given that mutations in the degenerate active site region decreased CspC levels (**Fig. 2B**), we tested whether the G171R mutation, which was isolated in 3 of the 10 germination mutants sequenced [12], would disrupt CspC folding. To this end, we recombinantly produced a G171R CspC variant in *E. coli* and compared its levels and purification yields to wild-type CspC and the CspC degenerate site mutants. Although CspC_G171R_-His_6_ was also produced to similar levels as wild-type CspC-His_6_ upon IPTG-induction (**Fig. S1B**, Induced), soluble CspC_G171R_-His_6_ purified with low efficiency relative to wild-type CspC-His_6,_ similar to the degenerate site mutants (**Fig. S1B**, Elution). These observations suggest that the loss-of-function CspC mutants identified in the germination mutant screen disrupt CspC folding and likely destabilize CspC in sporulating cells.

### The CspC prodomain cannot function in *trans*

Subtilisin-like serine proteases use their long N-terminal prodomain as an intramolecular chaperone to induce proper folding of the subtilase domain following translation [42]. Since the prodomains of other subtilisin-like serine proteases (including CspB in *C. difficile*) can perform this chaperone function *in trans* [33, 42], we tested whether the prodomain of *C. difficile* CspC could also function *in trans* even though it normally does not undergo autoprocessing. To this end, we generated a complementation construct that produced the prodomain (residues 1-64) separate from the remainder of CspC (residues 65-557, **Fig. 2A**). The resulting L64_TAG_ construct did not make detectable CspC in western blot analyses of sporulating cells or purified spores (**Fig. 2B**), suggesting that the CspC prodomain cannot function *in trans* unlike CspB and other subtilisin-like serine proteases.

Since we previously reported that the CspB prodomain could function *in trans* when over-expressed from a plasmid [32], we expressed this same construct (Q66_TAG_) in single-copy on the chromosome from the *pyrE* locus of a *cspBA* deletion strain (**Fig. S2A**). CspBA was detected in sporulating cells of the Q66_TAG_ complementation strain, albeit at reduced levels relative to wild type and the wild-type *cspBA* complementation strain (**Fig. S2B**). Thus, the CspB prodomain can act as a chaperone *in trans* to ensure proper folding of CspBA when *cspBA_Q66-TAG_* is expressed chromosomally, although with reduced efficiency. Consistent with our prior finding that CspB and CspA are important for incorporating and/or stabilizing CspC into mature spores [30, 38], the decreased CspB and CspA levels in purified Q66_TAG_ spores likely led to the reduced CspC levels observed (**Fig. S2B**).

Surprisingly, the greatly reduced levels of all three Csp proteins in Q66_TAG_ spores resulted in only a ~2-fold defect in spore germination on 0.1% taurocholate plates (**Fig. S2B**). Q66_TAG_ spores also exhibited wild-type levels of germinant sensitivity except for a ~4-fold decrease at the lowest concentration of taurocholate tested (0.001%, **Fig. S2C**). Thus, marked reductions in wild-type CspB, CspA, and CspC levels can mediate close to wild-type germination levels. Since CspBA_Q66-TAG_ spores, which have decreased levels of CspA, CspB, and CspC, and CspC_T170H_ spores, which only have decreased levels of CspC, exhibited similar germination responses (**Figs. 2** and **S2**), CspC would appear to be the primary determinant controlling germinant sensitivity.

### Structural flexibility of CspC appears to be important for function

The failure of the CspC prodomain to promote subtilase domain folding *in trans*, unlike CspB’s prodomain, may be a function of the higher number of interactions between CspC’s prodomain with both the subtilase and jelly roll domains. When the interfaces between each of the three subdomains were evaluated by buried surface area analysis, a larger contact area is observed between each of the domains in CspC as compared to CspB (**Table S1**, 3200 Å^2^ for CspC vs. 2270 Å^2^ for CspB, calculated with PDBe PISA[43]). Interestingly, despite the extensive interactions between the different sub-domains of CspC, residues in both the jelly roll domain and prodomain have higher B-factors than nearby residues in the subtilase domain (**Fig. 3A**). Since B-factors measure the movement of a residue around its average position, *C. difficile* CspC would appear to have greater flexibility in the jelly roll domain and prodomain than in the subtilase domain. In contrast, the previously solved crystal structure of *C. perfringens* CspB showed little flexibility in terms of its B-factors and was highly resistant to proteolysis [32]. CspC’s flexibility was particularly surprising given that the free energy of binding of the prodomain to the rest of the protein was −25.2 ΔG kcal/mol relative to −20.0 ΔG kcal/mol for CspB (**Table S1**). To determine if the high B-factors in *C. difficile* CspC result in high conformational flexibility relative to *C. perfringens* CspB, we subjected purified *C. difficile* CspC and *C. perfringens* CspB to limited proteolysis. Consistent with prior work, purified *C. perfringens* CspB was highly resistant to proteolysis by chymotrypsin, whereas purified *C. difficile* CspC was considerably more sensitive **(Fig. 3B**). These results indicate that *C. difficile* CspC is more conformationally dynamic than *C. perfringens* CspB.

**Figure 3.**
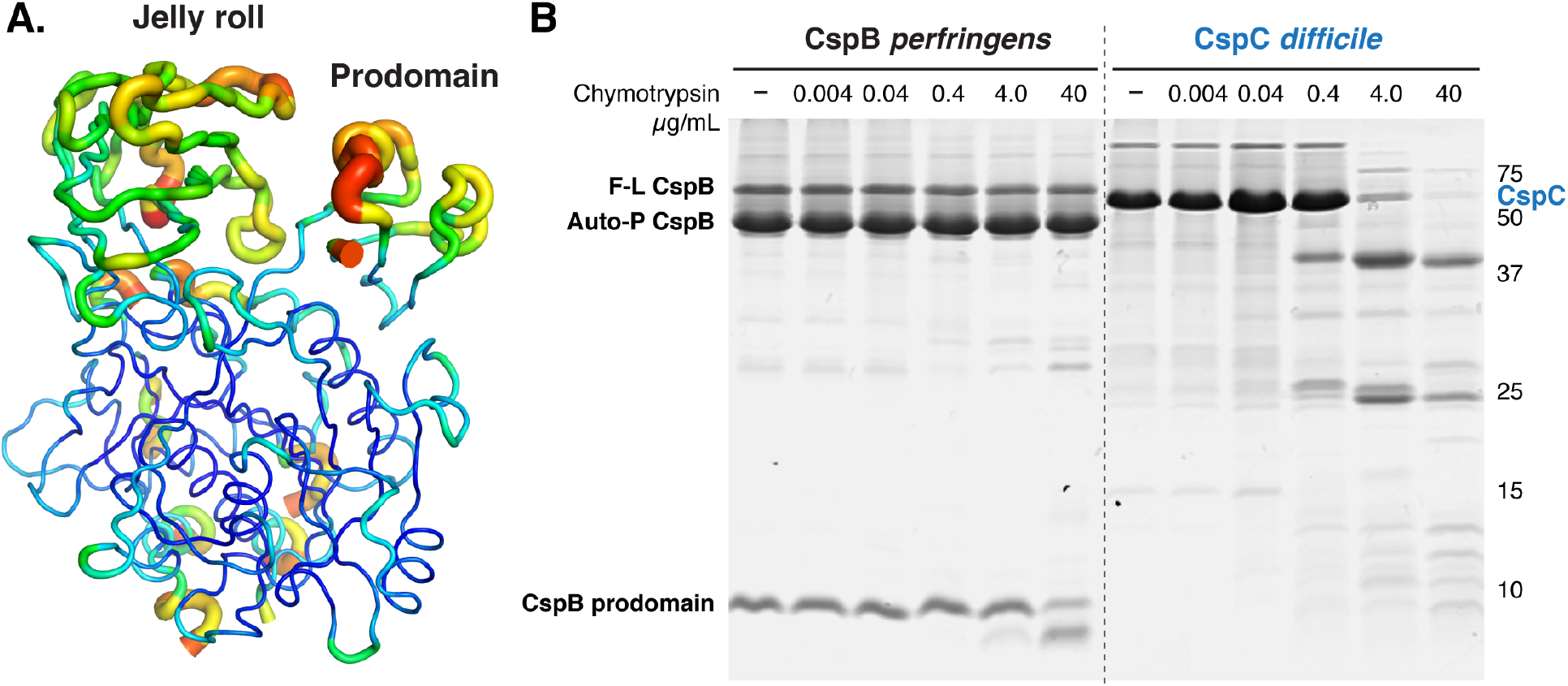
CspC is more conformationally flexible than CspB. (A) B-factor representation suggesting that the jelly roll and prodomains of CspC exhibit dynamic mobility. The higher relative B-factors are more striking for the jelly roll because there are strong crystal packing interfaces at two of the crystallographic two-folds. B-factor is indicated by color and line thickness. High B-factors represented by warm colors and a thick line; low B-factors represented by cool colors and a thin line. (B) Limited proteolysis of CspB and CspC. 15 μM of the proteins were incubated with increasing concentrations of chymotrypsin for 1 hr at 37°C. F-L refers to full-length CspB, and Auto-P refers to CspB that has undergone autoprocessing to release the CspB prodomain.

To assess whether the relative mobility of *C. difficile* CspC is important for its function, we targeted residues exhibiting conformational flexibility in the structure for mutagenesis. Both arginine 358 in the jelly roll and glutamate 43 in the prodomain have higher average B-factors (49.1 and 55.8 Å^2^, respectively, vs. an average of 27.2 Å^2^). These two residues form a salt bridge adjacent to another salt bridge between arginine 374 in the jelly roll and glutamate 57 in the prodomain, which have B-factors of 43.9 and 33.6 Å^2^, respectively (**Fig. 4A**). To test if the flexible Arg358 and Glu43 residues, or the neighboring salt bridge residues, are important for CspC function, we generated strains producing alanine substitutions of Glu43, Glu57, and Arg374, as well as alanine, glutamic acid, leucine substitutions at Arg358. None of these mutants exhibited statistically significant germination defects when plated on rich media containing 0.1% taurocholate (**Fig. 4B**). However, all the Arg358 mutants exhibited ~10-30-fold reduced sensitivity to germinants relative to wild-type when plated on media containing lower concentrations of germinant (i.e. 0.001% and 0.01% taurocholate, p < 0.005), whereas the remaining salt bridge mutant exhibited wild-type germinant sensitivity (**Fig. 4C**). Furthermore, the R358A mutant was severely impaired for germination in the optical density assay (**Fig. 4D**), indicating that it germinates more slowly than wild-type spores.

**Figure 4.**
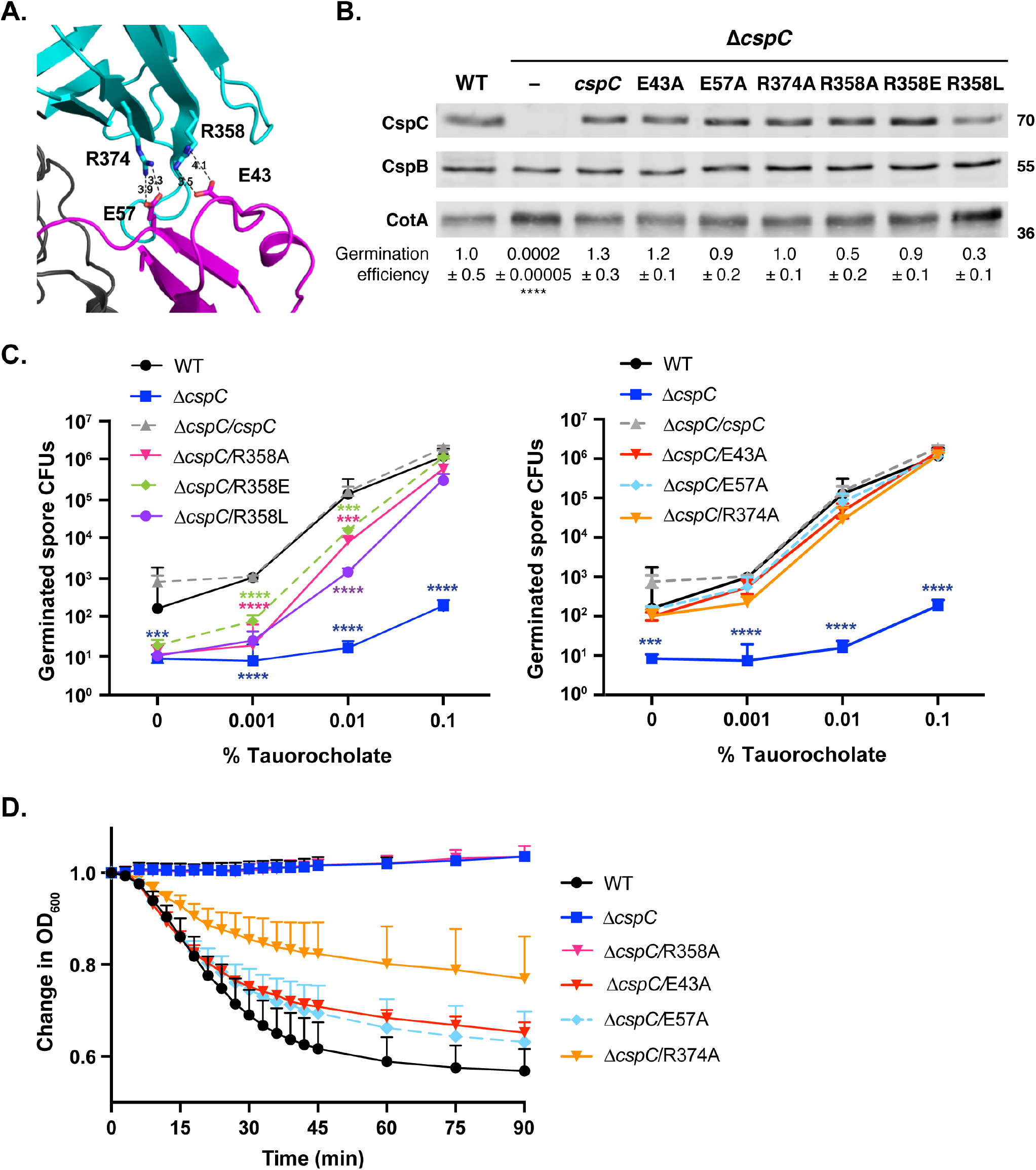
Mutations in the flexible residue, Arg358, decrease sensitivity to bile salt germinant. (A) Close-up view of dual salt-bridge interactions at the jellyroll domain-prodomain interface. The jelly roll domain is shown in cyan, and the prodomain is shown in magenta. The distances between the E57:R374 and R358:E43 salt bridge interactions are marked. (B) Western blot analyses of CspC and CspB levels in salt bridge mutant spores. CotA serves as a loading control. The germination efficiency of the salt bridge mutants plated on BHIS media containing 0.1% taurocholate is also shown relative to wild type. No significant differences in germination were observed for the salt bridge mutants using a one-way ANOVA and Tukey’s test. **** p < 0.0001. (C) Germinant sensitivity of salt bridge mutant spores plated on BHIS containing increasing concentrations of taurocholate. The number of colony forming units (CFUs) that arose from germinating spores is shown in two graphs to improve readability. For all germination assays shown, the mean and standard deviations shown are based on three biological replicates performed on two independent spore purifications. Statistical significance relative to wild type was determined using a one-way ANOVA and Tukey’s test. **** p < 0.0001, *** p < 0.001. (D) Optical density (OD_600_) analyses of spore germination over time in salt bridge mutants. Purified spores from the indicated strains were incubated in BHIS with and without 1% taurocholate, and the OD_600_ of the samples was monitored using a spectrophotometer. The change in OD_600_ represents the ratio of the OD_600_ of the treated to the untreated sample relative to the ratio at time zero. The averages of the results from three biological replicates performed on at least two independent spore preps is shown. The error bars indicate the standard deviation for each timepoint measured.

Importantly, none of the substitutions at Glu43, Glu57, Arg358, or Arg374 affected CspC levels in purified spores (**Fig. 4B**). Thus, the Arg358 substitutions specifically disrupt CspC function. Since mutations that flip or neutralize the charge on Arg358 affect germinant sensitivity, while mutation of Arg358’s salt bridge partner Glu43 (or the neighboring salt bridge) does not affect CspC function, the individual salt bridges do not appear to affect CspC function. These results suggest that the physical properties of Arg358, including potentially its conformational flexibility, are critical for CspC signaling.

### Mutation of the flexible residues G457 and R456 permits taurocholate-independent germination

Two additional residues exhibited considerable conformational flexibility in the CspC structure: arginine 456 and glycine 457, which lacked sufficient electron density to be included in our model for CspC (**Fig. 5A**). As described earlier, Francis *et al*. showed that a G457R mutation in CspC could expand the germinant specificity of *C. difficile* spores, enabling germination in response to chenodeoxycholate [12], which typically inhibits *C. difficile* spore germination [25]. This observation led to the hypothesis that CspC directly recognizes bile salts using glycine 457 [12].

Interestingly, Gly457 and Arg456 sit superior to a pocket formed on the surface of the subtilase domain that could potentially accommodate a molecule of taurocholate (data not shown). To test the importance of these residues, we individually mutated Gly457 to arginine and Arg456 to Gly. These mutations generate either two consecutive arginines or two consecutive glycines in the unstructured region, respectively. However, before characterizing the germinant sensitivity and specificity of these strains, we first tested if the G457R mutation in the 630Δ*erm* strain background would permit germination in response to chenodeoxycholate as previously described for a G457R mutant in the UK1 strain background [12]. To this end, we monitored the change in optical density of germinating spores in response to taurocholate and chenodeoxycholate in a plate reader as performed previously [12]. In our study, the *cspC*_G457R_ allele was expressed from the 630Δ*erm* chromosome, while the previous work expressed this mutant allele from a multicopy plasmid in strain UK1. CspC_G457R_ mutant spores germinated faster than wild-type spores in response to 10 mM taurocholate, suggesting that the glycine to arginine substitution at residue 457 increases the responsiveness of spores to taurocholate (**Fig. 5B**). Although CspC_G457R_ spores and wild-type spores decreased in optical density when exposed to 5 mM chenodeoxycholate, germination-defective Δ*cspC* spores also exhibited the same decrease in OD_600_ when exposed to chenodeoxycholate (**Fig. 5B**). These observations suggest that chenodeoxycholate causes non-specific effects on optical density in the tested conditions. Indeed, chenodeoxycholate exhibited low solubility in BHIS media at 37°C and formed a precipitate during the experiment.

**Figure 5.**
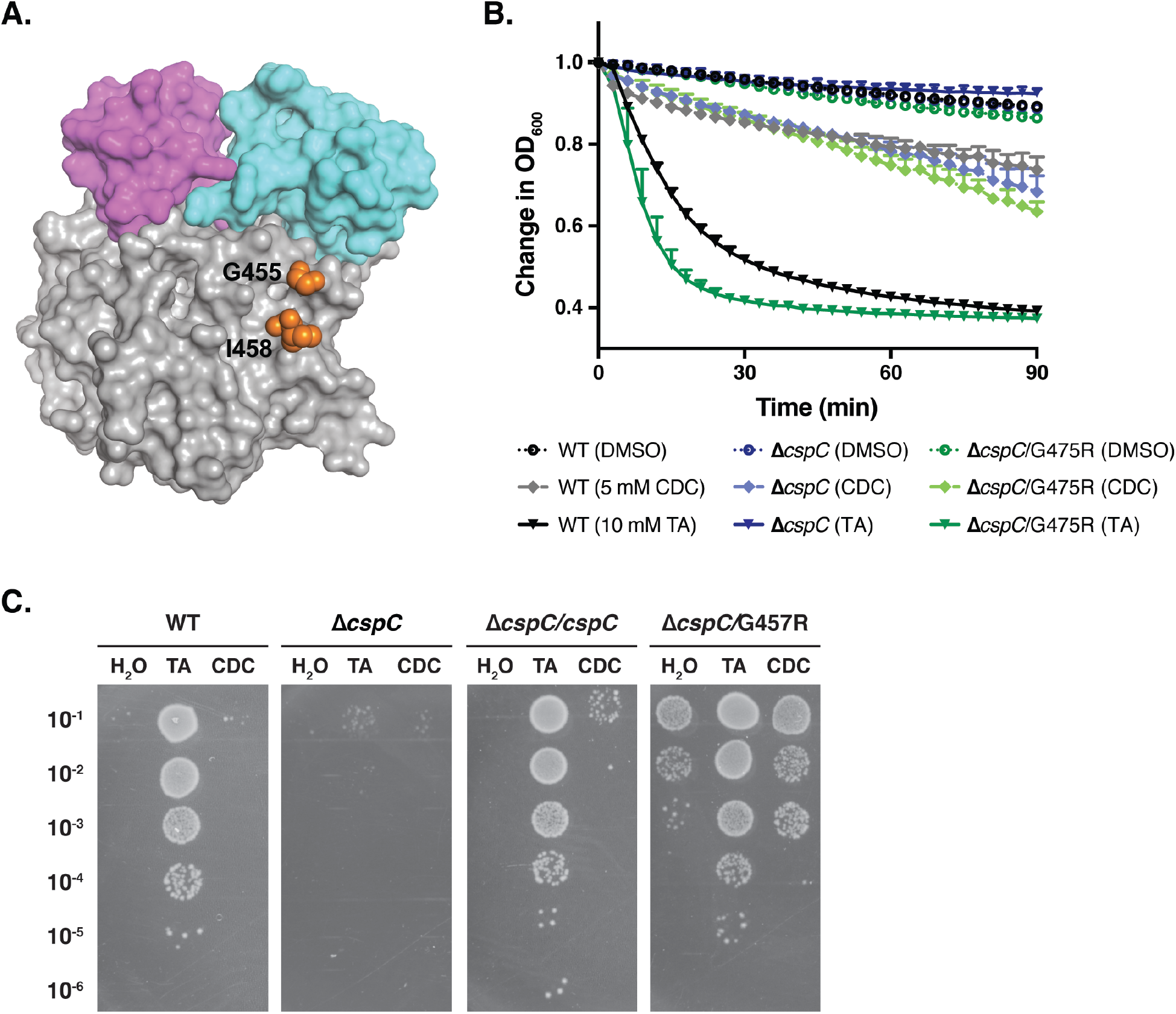
A G457R mutation results in taurocholate- and chenodeoxycholate-independent spore germination. (A) Space fill model of CspC highlighting the residues flanking (orange) the solvent-exposed Arg456 and Gly457 flexible loop. (B) Optical density (OD_600_) analyses of spore germination over time in the indicated strains. Purified spores from the indicated strains were incubated in BHIS in the presence of either DMSO carrier, 10 mM taurocholate, or 5 mM chenodeoxycholate. The OD_600_ of the samples was monitored in a 96-well plate using a plate reader. The change in OD_600_ represents the OD_600_ of the sample at a given timepoint relative to its starting OD_600_ at time zero. The averages of the results from three biological replicates are shown. The error bars indicate the standard deviation for each timepoint measured. (C) Purified spores from wild type, Δ*cspC*, and Δ*cspC* complemented with either wild-type CspC or a G457R mutant were incubated in BHIS supplemented with either water, 1% TA (19 mM) taurocholate (TA), or 0.5% (12 mM) chenodeoxycholate (CDC) for 30 minutes at 37°C then serially diluted in PBS and plated onto BHIS media lacking germinant. Colonies formed after a 24 hr incubation at 37°C are shown.than with taurocholate. Surprisingly, CspC_G457R_ spores also produced colonies when mock treated with water to the same extent as with chenodeoxycholate pretreatment (**Fig. 5C** & **S3**). These results indicate that, rather than altering germinant specificity, the *cspC*_G457R_ allele allows for germination *independent* of bile salt germinants.

To prevent chenodeoxycholate precipitation from confounding our germination measurements, we assessed the ability of CspC_G457R_ spores to germinate in response to pre-treatment with either taurocholate or chenodeoxycholate and form colonies when plated on media lacking germinant (BHIS). Colonies that form on BHIS alone arise from spores that germinated during the pre-treatment. Wild-type and wild-type complementation spores formed colonies only after pre-treatment with taurocholate and not chenodeoxycholate (**Fig. 5C & S3**). However, CspC_G457R_ mutant spores formed colonies in response to pre-treatment with both taurocholate and chenodeoxycholate, although ~100-fold less efficiently with chenodeoxycholate Since the rich BHIS media used in the experiments detailed above is poorly defined, we tested whether CspC_G457R_ spores would also germinate on a more minimal medium (*C. difficile* defined-media, CDDM, [44]). Pre-treatment of wild-type, Δ*cspC*, Δ*cspC*/*cspC*, and Δ*cspC*/G457R mutant spores with either water, taurocholate, or chenodeoxycholate followed by plating on CDDM alone yielded results nearly identical to that obtained on BHIS alone (**Fig. S3**). Thus, CspC_G457R_ mutant spores can germinate independently of a bile salt signal and appear to sense something present in both undefined rich media (BHIS) and minimal media (CDDM). Nevertheless, these mutant spores still respond to taurocholate as a germinant, since CFUs increase by ~2-logs upon taurocholate pre-treatment (**Figs. 5** and **S3**).

To test whether bile salt-independent germination would also be observed with the R456G substitution, we measured the germination sensitivity of CspC_R456G_ and CspC_G457R_ spores on BHIS lacking germinant and with increasing amounts of taurocholate. The R456G mutant spores also exhibited increased taurocholate-independent germination with a >1,000-fold increase in CFUs on BHIS alone relative to wild-type (**Fig. 6A**). CspC_R456G_ and CspC_G457R_ spore germination steadily increased in response to increasing taurocholate and reached maximum germination at 0.01% taurocholate, whereas wild-type and *ΔcspC/*cspC spore germination reached maximum germination at 0.1% taurocholate (**Fig. 6A**).

**Figure 6.**
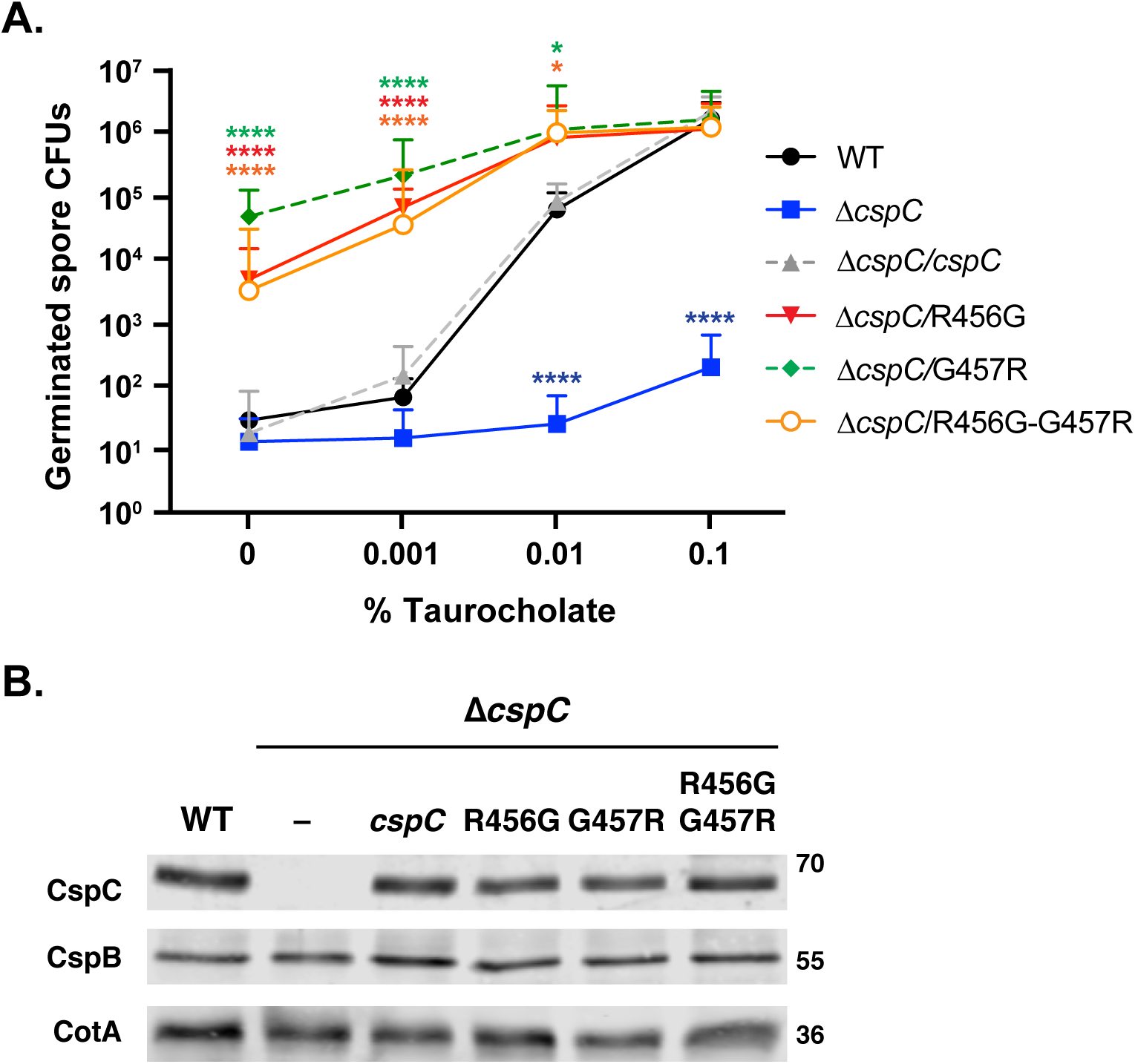
Mutations in the unstructured Arg456 and Gly457 result in increased taurocholate-independent germination. (A) Germinant sensitivity of *cspC* mutant spores encoding mutations in the flexible Arg456 and Gly457 when plated on BHIS containing increasing concentrations of taurocholate. The number of colony forming units (CFUs) produced by germinating spores is shown. The mean and standard deviations shown are based on four biological replicates performed on two independent spore purifications. Statistical significance relative to wild type was determined using a one-way ANOVA and Tukey’s test. **** p < 0.0001, *** p < 0.001. (B) Western blot analyses of CspC and CspB levels in G457 and R456 mutant spores. CotA serves as a loading control.

Since the G457R mutation generates an Arg-Arg stretch, and the R456G substitution makes a Gly-Gly stretch, we tested whether we could revert the taurocholate-independent germination phenotype back to wild-type levels by generating a R456G-G457R double mutant. This mutation reverses the order of amino acids at residues 456 and 457 relative to the wild-type protein.

Interestingly, CspC_R456G/G457R_ mutant spores exhibited the same phenotype as CspC_R456G_ mutant spores when plated on media with varying levels of taurocholate (**Fig. 6A**), forming colonies on media lacking germinant ~10-fold less efficiently than CspC_G457R_ spores, although this difference was not statistically significant. Importantly, all three mutants produced spores with wild-type amounts of CspC as determined by western blotting, suggesting that the CspC variants have enhanced signaling properties (**Fig. 6B**).

We next wondered what effect the identity of the residue at position 457 had on spore germination. To this end, we mutated Gly457 to another small, neutral amino acid (alanine), a polar residue (glutamine), a negatively charged residue (glutamic acid), and a smaller, positively charged amino acid (lysine). CspC_G457A_ and CspC_G457Q_ spores exhibited wild-type germination responses on BHIS alone and in response to different concentrations of taurocholate (**Fig. S4A**). Although slightly fewer CFUs were formed by these mutants compared to wild-type spores, this difference was not statistically significant. CspC_G457E_ spores exhibited an intermediate phenotype between that of wild-type and CspC_G457R_ spores, while CspC_G457K_ spores behaved identically to CspC_G457R_ spores (**Fig. S4A**). Western blot analysis indicated that none of these mutations affected CspC levels in mature spores (**Fig. S4B**). Taken together, charged amino acid substitutions of Gly456 (glutamic acid, lysine, and arginine) result in increased levels of taurocholate-independent germination, with positively charged amino acids having the greatest effect.

### Mutations of the flexible residues G457 and R456 increase sensitivity to taurocholate

Since high basal levels of germination in the absence of taurocholate made it difficult to distinguish in the plate-based taurocholate titration assay if the Arg456 and Gly457 mutations increase germinant sensitivity, we assessed their germinant sensitivity using the optical density-based germination assay. CspC_R456G_ and CspC_G457R_ spores were exposed to increasing concentrations of taurocholate in the presence of BHIS, and the decrease in optical density was measured compared to wild-type spores and Δ*cspC* spores. Both CspC_R456G_ and CspC_G457R_ spores germinated more quickly at all concentrations of taurocholate tested compared to wild-type spores (**Fig. S5**). The CspC_R456G_ and CspC_G457R_ spores in BHIS alone did not exhibit detectable germination in the optical density assay in contrast with the plate-based germination assays. This is presumably because the optical density assay lacks sufficient sensitivity to detect changes in <1% of the population (**Figs. 5, 6** and **S5**). Thus, in addition to having increased taurocholate-independent germination, the R456G and G457R variants have increased sensitivity to taurocholate germinant.

### A D429K mutation also leads to taurocholate-independent germination

The Arg456 and Gly457 residues are located on one edge of a pocket that could potentially bind small molecules, like taurocholate, so we tested if other residues in this region impact sensitivity to taurocholate. To this end, we made substitutions at residues on the other side of this pocket in aspartate 429 and glutamine 516. The substitutions either introduced a bulkier residue or reversed the charge of the native residue to potentially occlude ligand binding by this pocket (**Fig. 7A**). Of the four substitutions tested, only CspC_D429K_ exhibited a germination profile similar to CspC_R456G_ and CspC_G457R_ spores in that CspC_D429K_ spores germinated on BHIS alone at ~1000-fold higher levels than wild type (**Fig. 7B**, p < 0.005). Although germination saturated at 0.01% taurocholate like CspC_R456G_ and CspC_G457R_ spores, the shape of its germinant sensitivity curve resembled that of wild type in that its germination on 0.001% TA was only ~2-fold higher than in the absence of TA. In contrast, CspC_R456G_ and CspC_G457R_ spore germination increased by 15-and 5-fold, respectively, when comparing the germination levels on BHIS alone to BHIS containing 0.001% TA. Consistent with these observations, CspC_D429K_ spores did not exhibit greater sensitivity to taurocholate in the optical density assay than wild-type spores (**Fig.S5**), in contrast with CspC_R456G_ and CspC_G457R_ spores, indicating that the *D429K* allele results in a different germinant profile than the *R456G* and *G457R* alleles. Notably, none of the amino acid substitutions affected CspC levels in mature spores relative to wild-type (**Fig. 7C**).

**Figure 7.**
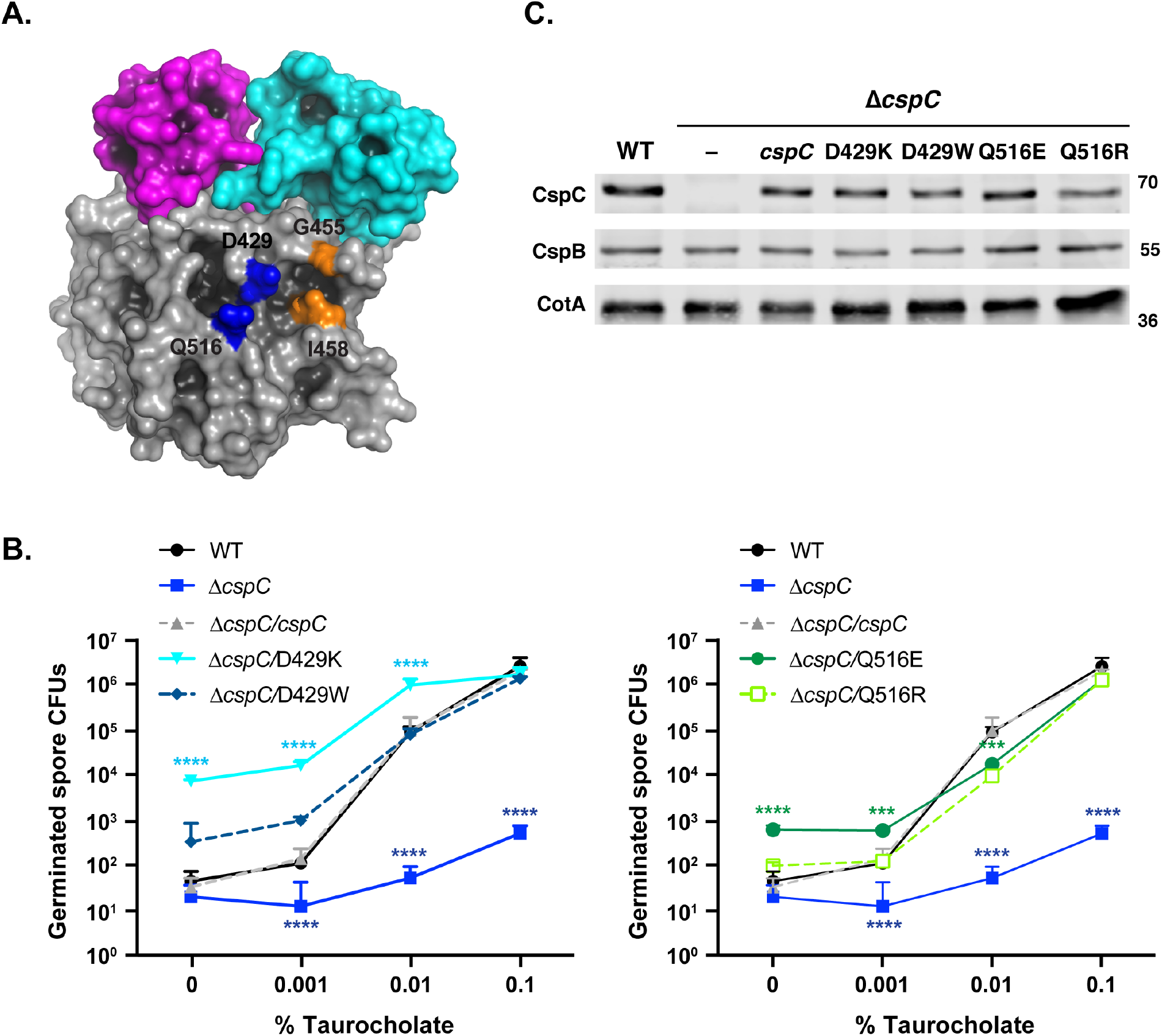
Mutation of Asp429 to Lys increases taurocholate-independent germination. (A) Space fill model of CspC highlighting the residues flanking the short loop containing R456 and G457 (orange spheres) juxtaposed to D429 and Q516 (blue spheres). A shallow cavity on the surface of the subtilase-like domain is visible between these four residues. The image is rendered to enhance cavity contrast using PyMOL (Schrödinger, Inc.) (B) Germinant sensitivity of *cspC* mutant spores encoding mutations in Asp429 and Gln516 when plated on BHIS containing increasing concentrations of taurocholate. The number of colony forming units (CFUs) that arose from germinating spores is shown in two graphs to improve readability. The mean and standard deviations shown are based on three biological replicates performed on two independent spore purifications. Statistical significance relative to wild type was determined using a one-way ANOVA and Tukey’s test. **** p < 0.0001, * p < 0.05. (B) Western blot analyses of CspC and CspB levels in D429 and Q516 mutant spores. CotA serves as a loading control.

### Mutants with increased taurocholate-independent germination are more sensitive to co-germinants

Although CspC_R456G_ and CspC_G457R_ spores differed from CspC_D429K_ spores in their sensitivity to taurocholate (**Fig. S5**), it was unclear why all three mutant spores germinated significantly better on plates in the absence of taurocholate germinant than wild-type spores. We hypothesized that the mutations confer differential responsiveness to components within both rich and defined media. Sorg and Sonenshein previously established that efficient *C. difficile* spore germination requires a second co-germinant signal [24], namely amino acids [45], which enhance sensitivity to taurocholate without causing germination on their own [19, 20]. Recently, Kochan *et al*. determined that divalent cations, particularly calcium, potentiate taurocholate-induced germination and that amino acid and calcium co-germinant signals can synergize to further enhance taurocholate-induced germination[46, 47]. It is currently unknown how *C. difficile* spores sense and respond to these co-germinants. Nevertheless, since amino acid and calcium co-germinants are present in the BHIS media used in our germination assays, we tested the sensitivity of CspC_R456G_, CspC_G457R_, and CspC_D429K_ spores to various co-germinants individually in the optical density assay using a PBS-based buffer rather than BHIS. We first tested the sensitivity of CspC_R456G_, CspC_G457R_, and CspC_D429K_ spores to the most potent amino acid co-germinant, glycine, and an amino acid co-germinant with a mid-range activating concentration (EC_50_), arginine [45]. Wild-type, CspC_R456G_, CspC_G457R_, and CspC_D429K_ spores did not germinate in PBS with 1% taurocholate alone (**Fig. 8**), just as BHIS alone did not induce germination in the optical density assay (**Fig. S5**). Thus, both germinant and a co-germinant must be present to detect germination in this assay. CspC_G457R_ spores germinated in lower concentrations of glycine (0.4 mM) than wild-type, CspC_R456G_, and CspC_D429K_ spores in the presence of 1% taurocholate and germinated faster than wild-type and CspC_D429K_ spores through all glycine concentrations tested (**Fig. 8**). At glycine concentrations of 2 mM and above CspC_R456G_ spores germinated faster than wild-type and CspC_D429K_ spores in the presence of 1% taurocholate (**Fig. 8A**). CspC_G457R_ and CspC_R456G_ spores also germinated faster than wild-type and CspC_D429K_ spores at 11.1 mM arginine with 1% taurocholate, with CspC_G457R_ spores germinating the fastest and at a lower arginine concentration (3.7 mM, **Fig. 8B**). Both CspC_R456G_ and CspC_G457R_ spores reached a maximum germination rate at 33.3 mM arginine with 1% taurocholate, while wild-type and CspC_D429K_ spores only reached similar germination rates at 100 mM arginine with 1% taurocholate (**Fig. 8B**). Taken together, these data indicate that CspC_R456G_ and CspC_G457R_ spores are more sensitive to glycine and arginine co-germinants in the presence of taurocholate, while CspC_D429K_ spores respond to these amino acid co-germinants similarly to wild-type spores.

**Figure 8.**
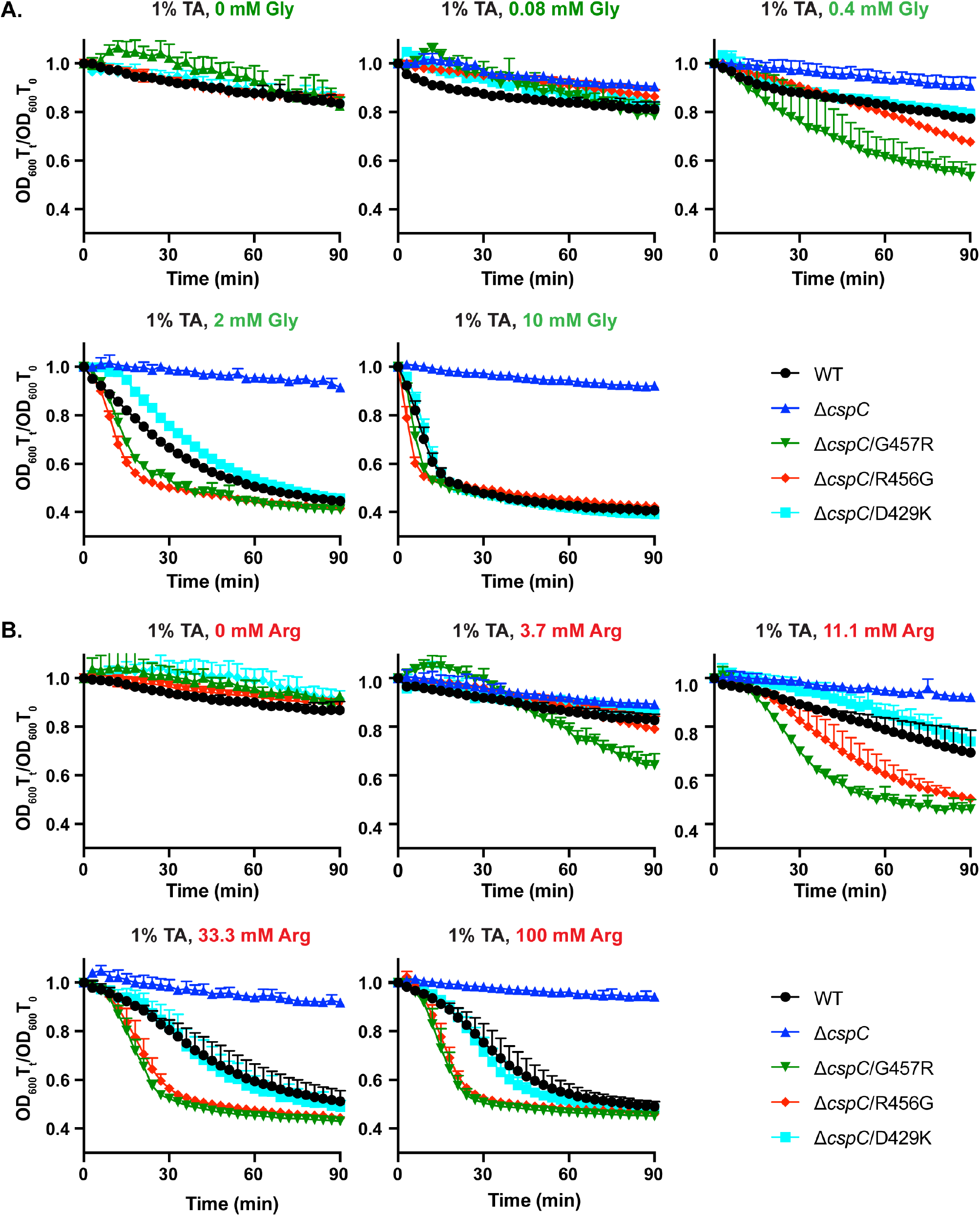
G457R and R456G mutations enhance sensitivity to amino acid co-germinants. (A) Optical density (OD_600_) analyses of spore germination over time in G457 region mutants. Purified spores from the indicated strains were incubated in PBS supplemented with 1% taurocholate and increasing concentrations of either (A) glycine or (B) arginine. The OD_600_ of the samples was monitored using a plate reader. The change in OD_600_ represents the OD_600_ of the sample at a given timepoint relative to its starting OD_600_ at time zero. The averages of the results from three biological replicates representative of least two independent spore preps are shown. The error bars indicate the standard deviation for each timepoint measured.

We next tested the sensitivity of these CspC mutants to calcium co-germinant. In PBS buffer with CaCl_2_ added, the spores clumped together, so we used Tris buffer to test the sensitivity of the CspC mutants to calcium. We also decreased the taurocholate concentration to 0.25% to determine the sensitivity of the spores to Ca^2+^, since spores germinated too rapidly in 1% taurocholate supplemented with CaCl_2_ to accurately measure the change in optical density in a plate reader. As observed with amino acid co-germinants, the optical density of all spores tested did not change in Tris buffer containing 0.25% taurocholate and no calcium (**Fig. 9**). However, CspC_G457R_ spores were highly sensitive to calcium, germinating at near maximal rates at the lowest calcium concentration tested (2.22 mM CaCl_2_, **Fig. 9**). CspC_D429K_ spores were also very sensitive to calcium, with the entire population germinating in response to 20 mM Ca^2+^. In contrast, wild-type and CspC_R456G_ spores did not germinate appreciably in response to 60 mM Ca^2+^ (**Fig. 9**), a result that differs slightly from that of Kochan *et al*., who first identified calcium as a co-germinant [47]. However, increasing the taurocholate levels to 1% in Tris buffer resulted in similar responses as Kochan *et al*. The different germinant sensitivities observed between the two studies could result from differences in the sporulation media (Clospore broth [47] vs. 70:30 plates) and spore purification methods. Regardless, our results indicate that CspC_R456G_, CspC_G457R_, and CspC_D429K_ spores exhibit differential sensitivity to one or more classes of co-germinants. This heightened sensitivity correlates with their increased taurocholate-independent germination on plates.

**Figure 9.**
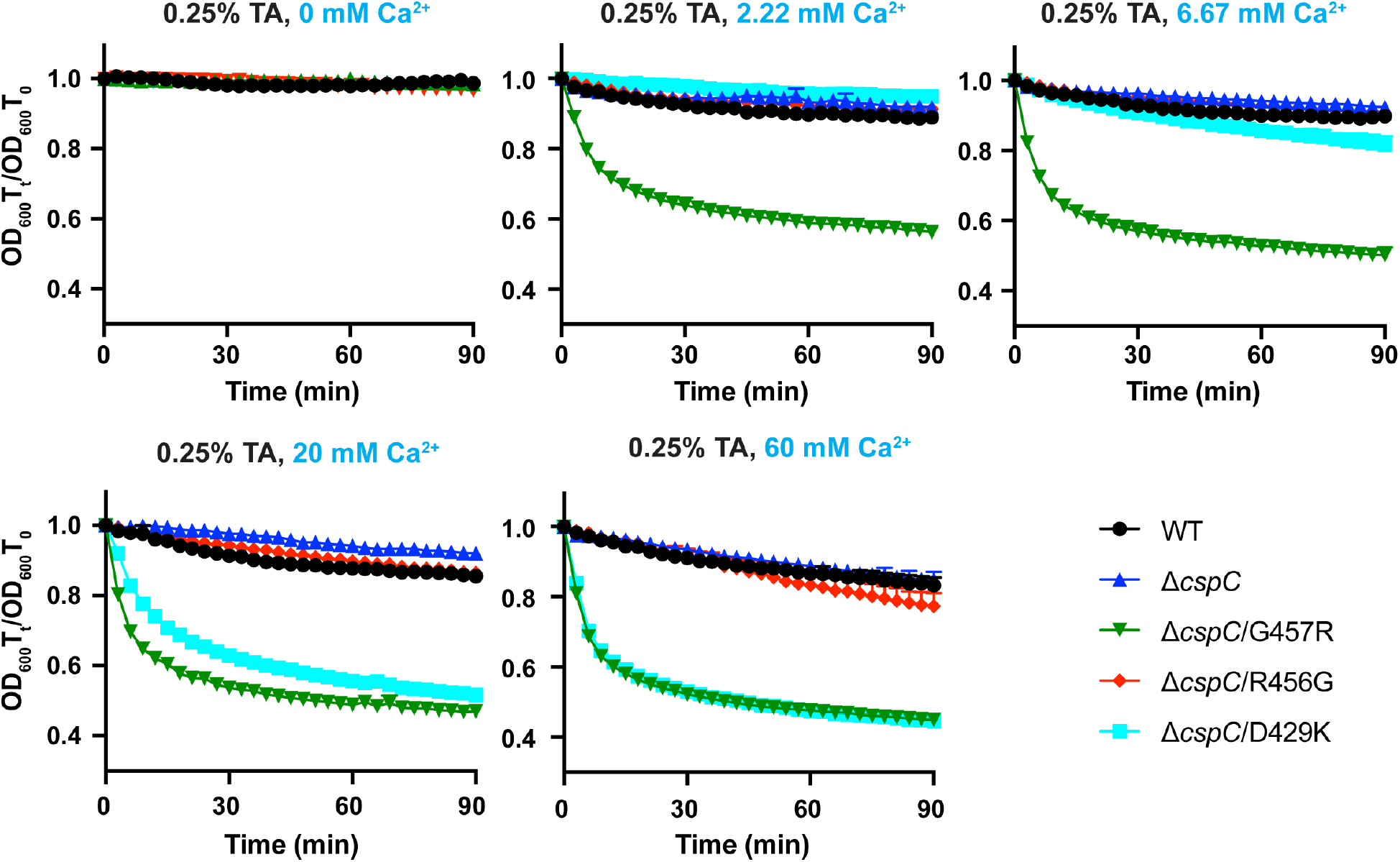
G457R and D429K mutations enhance sensitivity to the co-germinant, Ca^2+^. Optical density (OD_600_) analyses of spore germination over time in G457 region mutants. Purified spores from the indicated strains were incubated in Tris supplemented with 0.25% taurocholate and increasing concentrations of calcium. The OD_600_ of the samples was monitored using a plate reader. The change in OD_600_ represents the OD_600_ of the sample at a given timepoint relative to its starting OD_600_ at time zero. The averages of the results from three biological replicates representative of at least two independent spore preps are shown. The error bars indicate the standard deviation for each timepoint measured.

## Discussion

Unlike the nutrient germinants used by most spore-forming bacteria, the nosocomial pathogen, *C. difficile*, uses mammalian-specific bile salts as the signal for initiating germination. Furthermore, *C. difficile* lacks the membrane-bound germinant receptors common to almost all spore-forming bacteria [22, 48] and instead employs the Peptostreptococceae-specific CspC pseudoprotease to sense bile salt germinants [12]. By solving the X-ray crystal structure of CspC, we identified residues, particularly flexible residues, that are critical for CspC’s signaling function. Our mutational analyses surprisingly revealed that *C. difficile* CspC integrates signals not only from bile salt germinant but also from two different classes of co-germinants: amino acids and calcium (**Figs. 8** and **9**). Since the mechanism by which co-germinants are sensed by *C. difficile* was previously unknown, these observations provide important insight into how *C. difficile* spores integrate multiple environmental signals to induce germination. We also found that G457R mutant spores do not germinate in response to chenodeoxycholate (**Fig. 5, S3**), in contrast with a previous study [12]. Instead, the G457R mutation allows *C. difficile* spores to germinate on media in the absence of added bile salts. As we discuss later, these findings raise the possibility that *C. difficile* CspC may not bind bile salt germinants directly.

Our structure-guided mutational analyses also identified additional residues that regulate CspC function. These include the residues of CspC’s degenerate active site, since restoring the catalytic triad residues dramatically reduced CspC levels in sporulating *C. difficile* cells and did not activate autoprocessing in CspC (**Fig. 2**) even though CspC’s degenerate site residues and prodomain exhibit similar orientations as the CspB protease (**Fig. 1**). Since the mutations did not affect the production of these variants in *E. coli* but decreased their overall purification yield from soluble lysates (**Fig. S1**), CspC’s degenerate residues are critical for proper folding and/or stability of CspC. These findings are consistent with the strict conservation in degenerate site residues among CspC homologs in the Peptostreptococacceae family [30], although the role of CspC in these organisms remains unknown.

We also determined that conformationally flexible residues, Arg358, Arg456, and Gly457, are critical for CspC function, since mutation of these residues altered *C. difficile* CspC’s sensitivity to bile salt germinants (**Figs 4** and **S5**). Our finding that conformational mobility appears to be important for CspC function was somewhat surprising given that a unique hinge and clamp region in CspC provides a larger contact surface area between the prodomain and subtilase domain than in CspB (**Fig. 1A** and **Table S1**). Despite this larger contact surface, CspC has greater conformational flexibility than CspB as determined by limited proteolysis analyses (**Fig. 3**).

Mutation of one of these conformationally flexible residues, Arg358, in the jelly roll domain to chemically distinct residues (alanine, glutamatic acid, and leucine) resulted in hypo-sensitivity to taurocholate germinant and a dramatically reduced germination rate (**Fig. 4**). Although Arg358 can form a salt bridge with the nearby Glu43, this interaction appears dispensable for CspC function, since an E43A mutation did not affect germination rate or germinant sensitivity (**Fig. 4**). Thus, Arg358’s ability to interact with the surrounding environment likely determines CspC’s ability to sense germinant and/or transduce the germinant signal.

The flexible residues, Arg456 and Gly457, are located almost 180° from Arg358 and the degenerate active site. Our results indicate that these residues are important for sensing and/or responding to bile salts and amino acids, since mutations of these residues surprisingly led to bile salt-independent spore germination (**Figs. 5** and **6**) as well as increased sensitivity to taurocholate germinant (**Fig. S5**) and amino acid co-germinants (**Fig. 8**). Mutations of Gly457 to charged residues in particular potentiated bile salt-independent spore germination (**Fig. S4**), suggesting that charged residues may promote interactions of CspC with (co)-germinants and/or signal transduction proteins.

We also identified D429K as another mutation that confers bile salt-independent germination to *C. difficile* spores. Asp429 borders a shallow pocket next to the predicted positions of R456 and G457 (**Fig. 7**); however, unlike the R456G and G457R mutations, the D429K mutation did not increase responsiveness to taurocholate or amino acids and instead markedly increased spore germination in response to the calcium ion co-germinant (**Fig. S5, 7-9**). In contrast, the R456G mutant exhibited wild-type responsiveness to Ca^2+^, while the G457R mutant was even more sensitive to Ca^2+^ than the D429K mutant (**Fig. 9**). Taken together, the D429K mutation increases sensitivity to calcium co-germinant alone; the R456G mutation increases sensitivity to taurocholate germinant and amino acid co-germinants; and the G457R mutation exhibits the highest sensitivity to all three of these small molecule types (**Figs, S5, 8**, **9**).

Interestingly, of the mutants we characterized, only those that were more sensitive to amino acid co-germinants were more sensitive to taurocholate germinant. Amino acid co-germinants were recently reported to act synergistically with divalent cation co-germinants, but neither class of co-germinants synergizes with members of the same class [46]. The authors hypothesized that amino acid co-germinants only synergize with divalent cation co-germinants because the different classes of co-germinants are sensed by different receptors or potentially even different pathways [20, 46]. Our current data do not distinguish between whether *C. difficile* spores use two separate receptors, the same receptor, or CspC itself, to sense amino acid and divalent cations, but it is interesting that the G457R mutation increases sensitivity to both classes of co-germinants and that the Gly457, Arg456, and D429 residues all line the edge of a pocket that could bind taurocholate and/or small molecules like amino acids.

While CspC clearly integrates multiple small molecule signaling inputs, the mechanism by which it senses and responds to these chemically distinct classes of molecules (bile salts, amino acids, and divalent cations) is unknown. CspC may bind bile salts, amino acids, and divalent cations directly or indirectly. While Ca^2+^ is a co-factor in some subtilisin-like proteases [33, 49], we did not observe evidence for Ca^2+^ binding in the structure. If CspC indirectly senses these molecules, it presumably interacts with the direct receptors for bile salts and co-germinants. In this scenario, the mutations we identified would enhance CspC’s association with these receptors. Alternatively, CspC could function through some combination of these models by directly binding germinant and a co-germinant receptor(s) or vice versa. A final model that has been proposed is that CspC alters the permeability of germinants to their “true” receptors in the cortex or core [20]. It is currently unknown whether bile salt germinants and Ca^2+^ co-germinants can pass through the outer forespore membrane to reach the cortex where CspC and CspB are hypothesized to be located [19, 28, 35]. Thus, CspC may facilitate the transport of germinants and/or co-germinants to the cortex region. Clearly, more work needs to be done to distinguish between these models.

Our findings also raise the question as to how CspC activation by taurocholate and co-germinants (directly or indirectly) leads to activation of the CspB protease. CspC could activate CspB through a direct interaction [19, 35, 38], as some subtilisin-like proteases form dimers [49, 50]. This mechanism would be consistent with the observation that allosteric interactions between pseudoenyzmes can activate their cognate enzymes [51-56]. A possible model for CspB activation by CspC is that direct or indirect binding of germinants and co-germinants induces a conformational change in CspC that allows CspC to bring CspB into an active conformation through a direct interaction. Our observation that CspC is conformationally flexible may reflect the need for CspC to adopt a different conformation in response to germinant and/or co-germinant signal(s) that then allows CspC to activate CspB. Further work will need to be done to test these hypotheses.

In addition to providing insight into the function of CspC, the mutations we have identified could help answer one of the major questions in the field regarding how sensitivity to germinant affects the ability of *C. difficile* to cause disease. Epidemic strains of *C. difficile* vary in the severity of the disease they cause and their sensitivity to bile salt germinant [57-61]. However, since *C. difficile* strains exhibit high genetic diversity, it has not been possible to directly correlate germinant sensitivity to disease severity. High sensitivity to bile salt germinants could increase the effective infectious dose of *C. difficile* and thus lead to more severe disease, or lower sensitivity to germinant could ensure that *C. difficile* spores germinate closer to the colon, the primary site of infection, and thus cause more severe disease [57, 60-62]. Testing the ability of our hypo- and hyper-sensitive mutants to colonize and cause disease in animal infection models could help answer this outstanding question in the field about *C. difficile* pathogenesis.

## Materials and Methods

### Bacterial strains and growth conditions

All *C. difficile* strain manipulations were performed with 630Δ*ermΔpyrE* as the parental strain using *pyrE*-based allele-coupled exchange (ACE [37]). *C. difficile* strains are listed in **Table S2**; they were grown on brain heart infusion media (BHIS) supplemented with L-cysteine (0.1% w/v; 8.25 mM), taurocholate (TA, 0.1% w/v; 1.9 mM), thiamphenicol (5-15 μg/mL), kanamycin (50 μg/mL), cefoxitin (8 μg/mL), as needed. Cultures were grown at 37°C under anaerobic conditions using a gas mixture containing 85% N_2_, 5% CO_2_, and 10% H_2_.

*Escherichia coli* strains for HB101/pRK24-based conjugations and BL21(DE3)-based protein production are listed in **Table S2**. *E. coli* strains were grown at 37 °C, shaking at 225 rpm in Luria-Bertani broth (LB). The media was supplemented with chloramphenicol (20 μg/mL), ampicillin (50 μg/mL), or kanamycin (30 μg/mL) as indicated.

### *E. coli* strain construction

Plasmid constructs were cloned into DH5α and sequence confirmed using Genewiz. To construct the *cspC* mutant complementation constructs, flanking primers, #2189 and 2242 (**Table S3**), were used in combination with internal primers encoding a given point mutation with Δ*cspBA* genomic DNA template. The resulting *cspC* complementation constructs carry 282 bp of the *cspBA* upstream region along with the Δ*cspBA* sequence and the intergenic region between *cspBA* and *cspC*. This slightly extended construct was necessary to generate wild-type levels of CspC when the constructs were expressed from the *pyrE* locus as previously described [38]. The primers encoding each CspC point mutation are provided in **Table S3**, where all primers used are listed.

For example, the T170H mutation was constructed using primer pair #2189 and 2355 to amplify a 5’ *cspC* complementation construct fragment encoding the T170H mutation at the 3’ end, while primer pair #2354 and 2242 were used to amplify a 3’ *cspC* complementation construct encoding the T170H mutation at the 5’ end. The individual 5’ and 3’products were cloned into pMTL-YN1C digested with NotI/XhoI by Gibson assembly. In some cases, the two PCR products were used in a PCR SOE [63] prior to using Gibson assembly to clone the *cspC* construct into pMTL-YN1C digested with NotI and XhoI. The resulting plasmids were transformed into *E. coli* DH5α, confirmed by sequencing, and transformed into HB101/pRK24.

To clone the construct encoding the *cspBA* prodomain trans-complementation construct, primer pair #2189 and 951 was used to amplify the 5’ fragment, and primer pair #950 and 2242 were used to amplify the 3’ fragment. In both cases, Δ*cspC* genomic DNA was used as a template, since it was necessary to include the downstream Δ*cspC* sequence to generate wild-type levels of CspBA in a wild-type complementation construct [38]. The resulting two fragments were joined together using PCR SOE with primer pair #2189 and 2242, and the PCR SOE product was cloned into pMTL-YN1C digested with NotI and XhoI using Gibson assembly. A similar strategy was used to generate the *cspC* prodomain trans-complementation construct. Primers #2189 and 2553 were used to amplify the 5’ fragment and primer pair #2552 and 2242 were used to amplify the 3’ fragment using Δ*cspBA* genomic DNA as a template. The fragments were joined together using SOE PCR with the primer pair #2189 and 2242, and the resulting SOE PCR product was cloned into pMTL-YN1C digested with NotI and XhoI using Gibson assembly.

To generate the recombinant protein expression constructs for producing CspC-His_6_ variants, primer pair #1128 and 1129 was used to amplify a codon-optimized version of *cspC* using pJS148 as the template (a kind gift from Joseph Sorg). The resulting PCR product was digested with NdeI and XhoI and ligated into pET22b cut with the same enzymes. The ligation mixture was used to transform DH5α. The G457R variant was cloned using a similar procedure except that primer pair #1128 and 1361 and primer pair #1360 and 1129 were used to PCR the 5’ and 3’ fragments encoding the G457R mutation. The resulting PCR products were joined together using PCR SOE and flanking primer pair #1128 and 1129.

The remaining constructs encoding *cspC* codon-optimized variants for expression using pET22b were cloned using Gibson assembly. Flanking primer pair #2311 and 2312 were used to generate PCR products when used in combination with the internal primers encoding the point mutations. The resulting PCR products were cloned into pET22b digested with NdeI and XhoI using Gibson assembly. PCR SOE was sometimes used to join the two 5’ and 3’ fragments prior to Gibson assembly into pET22b.

### Protein purification for crystallography

*E. coli* BL21(DE3) strains 981 and 1721 (**Table S2**) were used to produce codon-optimized CspC (wild-type and G457R, respectively) using the autoinduction method. Briefly, the indicated strains were struck out onto LB plates containing ampicillin and used to inoculate a 20 mL culture of LB containing 100 μg/mL ampicillin. The culture was grown for ~4 hrs after which it was used to inoculate 1:1000 of Terrific Broth (Affymetrix) supplemented with 5052 sugar mix (0.5% glycerol, 0.05% glucose, 0.1% alpha-lactose) and ampicillin for 60 hrs at 20°C [64]. The cells were pelleted and then resuspended in lysis buffer (500 mM NaCl, 50 mM Tris [pH 7.5], 15 mM imidazole, 10% [vol/vol] glycerol), flash frozen in liquid nitrogen, thawed and then sonicated. The insoluble material was pelleted, and the soluble fraction was incubated with Ni-NTA agarose beads (5 Prime) for 3 hrs, and eluted using high-imidazole buffer (500 mM NaCl, 50 mM Tris [pH 7.5], 200 mM imidazole, 10% [vol/vol] glycerol) after nutating the sample for 5-10 min.

Four elution fractions were pooled and then buffer exchanged into gel filtration buffer (200 mM NaCl, 10 mM Tris pH 7.5, 5% [vol/vol] glycerol) using Amicon 30 kDa cut-off filters. The buffer-exchanged protein was concentrated to 20 mg/mL or less, and gel filtration chromatography was performed using a Superdex 200 (GE Healthcare) column. Fractions containing CspC-His6 were pooled, concentrated to ~30 mg/mL, and flash frozen in liquid nitrogen.

### Crystallization and structure determination

Purified protein was buffer exchanged into 25 mM Hepes pH 7.5, 100 mM NaCl, 1 mM TCEP with 10 % [vol/vol] glycerol and concentrated to 22 mg/ml. Crystallization was performed via hanging drop vapor diffusion method by mixing of equivalent volumes of protein to crystallization reagent solution. Crystals grew in a broad range of conditions but suffered from severe twinning. This challenge was addressed by pre-incubation of the protein with 0.5 M guanidinium hydrochloride prior to mixing with crystallization reagent. Crystals grew with a reagent containing 0.2 M MES pH 6.5 and 2.5 M ammonium chloride equilibrating over a well containing 2M NaCl and incubated at room temperature. Cryoprotection was achieved by serial transfer from the growth condition to a final solution containing 1.0 M ammonium sulfate and 2.0 M lithium sulfate prior to cryo-cooling into liquid nitrogen.

Data were collected at 12 KeV on GM/CA beamline 23-ID-D at the Advanced Photon Source (Argonne National Laboratory) using a Pilatus3 6M detector with data processed to 1.55 Å using HKL2000 [65] (**Table 1**). The crystals were of space group C222(1) with a single molecule in the asymmetric unit. A molecular replacement search model was prepared from the subtilase-like domain of CspB (4I0W) [32] using a sequence alignment imported into Chainsaw [66] followed by truncation to a c-alpha only trace. A clear molecular replacement solution (TFZ of 10.3) was achieved using Phaser [67] within Phenix [32] and submitted to density modification using Solve/Resolve [68]. Clear density for both the prodomain and jelly roll domains was observed with model completion performed by iterative rounds of manual building, automated building using Autobuild and final refinement using Phenix with a final R_free_ of 18.6. The following residues were not included in the model due to ambiguous electron density: 1, 89-92, 456-457, 506-508, 556-558.

**Table 1.** Crystallographic data collection and refinement statistics.

|  |  |
| --- | --- |
| PDB ID Code | 6MW4 |
| Space Group | C222(1) |
| Cell a, b, c (Å) | 88.6, 155.2, 91.7 |
| Resolution Å | 50 – 1.55 |
| <sup>1</sup>(High resolution Å) | (1.61 – 1.55) |
| Reflections | 91876 |
| Completeness % | 99.9 (99.9) |
| Multiplicity | 6.5 (6.0) |
| Wilson B | 18.1 |
| I/SigmaI | 24.2 (1.7) |
| Rmeas % | 6.6 (17.8) |
| Rpim % | 2.6 (28.8) |
| CC 1/2 | 0.996 (0.857) |
| Coordinate error Å | 0.18 |
| Phase error ° | 18.1 |
| R<sub>work</sub> % | 16.4 |
| <sup>2</sup>R<sub>free</sub> % | 18.6 |
| Rms bonds Å | 0.006 |
| Rms angles ° | 0.806 |
| Ramachandran |  |
| Favored | 98.7 |
| Disallowed | 0.37 |
| Mean B Å<sup>2</sup> |  |
| Protein (atoms) | 27.1 (4214) |
| Solvent (atoms) | 36.9 (375) |
| Sulfate (atoms) | 56.4 (25) |

<sup>1</sup>Numbers in parentheses denote high resolution.
<sup>2</sup>Represents 10% of reflections omitted from refinement.

### Protein purification for His_6_-tag pulldown assays

*E. coli* BL21(DE3) strains with pET22b plasmids encoding codon-optimized variants of *cspC* (the wild-type gene was a kind gift of Joseph Sorg) (see **Table S2** in the supplemental material) were grown for protein purification to mid-log phase in 2YT (5 g NaCl, 10 g yeast extract, and 15 g tryptone per liter). When cultures reached an OD_600_ ~0.8, 250 μM isopropyl-β-D-1-thiogalactopyranoside (IPTG) was added to induce expression of *cspC*. Cultures were then grown overnight at 19°C after which they were pelleted, resuspended in lysis buffer (500 mM NaCl, 50 mM Tris [pH 7.5], 15 mM imidazole, 10% [vol/vol] glycerol), flash frozen in liquid nitrogen, thawed and then sonicated. The insoluble material was pelleted, and the soluble fraction was incubated with Ni-NTA agarose beads (0.5 ml; 5 Prime) for 3 hrs, and eluted using high-imidazole buffer (500 mM NaCl, 50 mM Tris [pH 7.5], 200 mM imidazole, 10% [vol/vol] glycerol) after nutating the sample for 5-10 min. Samples were removed prior to harvesting cells (input), after lysis and centrifugation (cleared lysate), and after elution for western blot analyses. A sample from the insoluble pellet was also analyzed after resuspending in 9M urea.

### Limited proteolysis

Purified *C. perfringens* CspB or *C. difficile* CspC proteins were diluted to 15 μM in 10 mM Tris pH 7.5 buffer. The protein solution was aliquoted into 0.2 mL tubes. A 1 mg/mL solution of chymotrypsin in 1 mM HCl was serially diluted to generate 10-fold dilutions for use in the assay. The serially diluted chymotrypsin solutions were added to the aliquoted protein solutions for a final concentration of chymotrypsin ranging from 40 μg/mL to 0.0004 μg/mL. 1 mM HCl was added to protein samples as an untreated control. The protein and chymotrypsin mixture was incubated at 37°C for 1 hr. Chymotrypsin activity was quenched by the addition of NuPAGE 4X LDS Sample Buffer (Invitrogen) and boiled at 98°C for three minutes. 10 μL aliquots were resolved using a 15% SDS-PAGE gel and visualized by Coomassie staining.

### *C. difficile* strain construction

Complementation strains were constructed as previously described using CDDM to select for recombination of the complementation construct into the *pyrE* locus by restoring uracil prototrophy [41]. Two independent clones from each complementation strain were phenotypically characterized.

### Sporulation

*C. difficile* strains inoculated from glycerol stocks were grown overnight on BHIS plates containing taurocholate (TA, 0.1% w/v, 1.9 mM). The resulting colonies were used to inoculate liquid BHIS cultures, which were grown to early stationary phase before being back-diluted 1:50 into BHIS. When the cultures reached an OD_600_ between 0.35 and 0.75, 120 μL of this culture were spread onto 70:30 agar plates and sporulation was induced as previously described [69] for 21-24 hrs. Sporulating cells were harvested into phosphate-buffered saline (PBS), and cells were visualized by phase-contrast microscopy.

### Spore purification

After inducing sporulation on 70:30 agar plates for 2-3 days as previously described [70], the samples were harvested into ice-cold, sterile water. A sample was removed to examine the sporulation cultures using phase-contrast microscopy. The spore samples were then washed 6 times in ice-cold water and incubated overnight in water at 4°C. The following day, the samples were pelleted and treated with DNase I (New England Biolabs) at 37 °C for 30-60 minutes, and purified on a 20%/50% HistoDenz (Sigma Aldrich) gradient. The resulting spores were washed 2-3 times in water, and spore purity was assessed using phase-contrast microscopy (>95% pure). The optical density of the spore stock was measured at OD_600_, and spores were stored in water at 4 °C.

### Germination assay

Germination assays were performed as previously described [71]. For each strain tested, the equivalent of 0.35 OD_600_ units, which corresponds to ~1 x 10^7^ spores, were resuspended in 100 μl of water, and 10 μL of this mixture were removed for 10-fold serial dilutions in PBS. The dilutions were plated on BHIS-TA, and colonies arising from germinated spores were enumerated at 20-24 hrs. Germination efficiencies were calculated by averaging the CFUs produced by spores for a given strain relative to the number produced by wild type spores for at least three biological replicates using spores from two independent preparations. Statistical significance was determined by performing a one-way analysis of variance (ANOVA) on natural log-transformed data using Tukey’s test. The data were transformed because the use of two independent spore preparations resulted in a non-normal distribution.

Germination assays with chenodeoxycholate or taurocholate pretreatment were performed essentially as previously described [30]. Briefly, ~3 x 10^7^ spores (1.05 OD_600_ units) were resuspended in 150 μL water. 150 μL of BHIS was added to the spore suspensions. Aliquots of the spore suspensions were exposed to 1% taurocholate, 5% chenodeoxycholate, or water (untreated) and incubated at 37°C for 20 minutes. 10 μL of this mixture was removed for 10-fold serial dilutions in PBS and the dilutions were plated on BHIS and BHIS with 0.1% taurocholate or CDDM and CDDM with 0.1% taurocholate. Colonies arising from germinated spores were enumerated at 20-24 hrs. Data presented are the averages of CFUs enumerated from at least three replicates performed on two independent spore preparations. Statistical significance was determined by performing a one-way analysis of variance (ANOVA) on natural log-transformed data using Tukey’s test. The data were transformed because the use of two independent spore preparations resulted in a non-normal distribution.

### OD_600_ kinetics assay – cuvette-based

Germination was induced as previously described [41]. About ~1.4 x 10^7^ spores (0.48 OD_600_ units) were re-suspended in BHIS and then exposed to 1% taurocholate (19 mM) or water (untreated sample). The OD_600_ of the TA- and untreated samples was measured every 3 min for 45 min, followed by every 15 min for a total of 90-120 min. The change in OD_600_ over time was determined by calculating the ratio of the OD_600_ measured for TA-treated samples to that for untreated samples relative to the ratio determined at time zero.

### OD_600_ kinetics assay – plate-based

For OD_600_ kinetics assays with chenodeoxycholate in 96-well plates, ~1.6 x 10^7^ spores (0.55 OD_600_ units) for each condition tested were resuspended in BHIS and 180 μL were aliquoted into three wells of a 96 well flat bottom tissue culture plate (Falcon) for each condition tested. The spores were exposed to 10 mM taurocholate, 5 mM chenodeoxycholate, or 50% DMSO (untreated) in a final volume of 200 μL. The OD_600_ of the samples was measured every 3 minutes in a Synergy H1 microplate reader (Biotek) at 37°C with constant shaking between readings. The OD_600_ for each technical triplicate was averaged at each time point and the OD_600_ of a blank measurement (BHIS with 0 mM taurocholate, 5 mM chenodeoxycholate, or 50% DMSO alone) was subtracted from the OD_600_ of the appropriately treated spores at each time point. The change in OD_600_ over time was calculated as the ratio of the OD_600_ at each time point to the OD_600_ at time zero.

For OD_600_ kinetics assays with varying concentrations of taurocholate germinant, ~2.3 x 10^7^ spores (0.8 OD_600_ units) for each condition tested were resuspended in BHIS and 900 μL were aliquoted into a well of a 24 well suspension culture plate (CellStar) for each condition tested. The spores were then exposed to 2-fold dilutions of 1% taurocholate (19 mM) or water (untreated) in a total volume of 1 mL. The OD_600_ of the samples was measured every 3 minutes in a Synergy H1 microplate reader (Biotek) at 37°C with constant shaking between readings. The change in OD_600_ over time was calculated as the ratio of the OD_600_ at each time point to the OD_600_ at time zero.

For OD_600_ kinetics assays with varying concentrations of co-germinants, ~2.3 x 10^7^ spores (0.8 OD_600_ units) for each condition tested were resuspended in either 1.5X PBS buffer or 50 mM Tris HCl pH 7.5 and aliquoted into a well of a 24-well plate for each condition tested. As spores clumped when CaCl_2_ was added to spores resuspended in 1.5X PBS, 50 mM Tris HCl pH 7.5 was used to measure OD600 kinetics in response to calcium as previously described [47]. 5-fold serial dilutions of 1 M glycine, 3-fold serial dilutions of 1 M arginine, 3-fold serial dilutions of 6 M CaCl_2_, or 1.5X PBS or 50 mM Tris HCl (untreated) were added to spores resuspended in the appropriate buffer to a final volume of 900 μL. The spores were then exposed to 1% taurocholate (19 mM) in a total volume of 1 mL and the OD_600_ of the samples was measured every 3 minutes in a Synergy H1 microplate reader (Biotek) at 37°C with constant shaking between readings. The change in OD_600_ over time was calculated as the ratio of the OD_600_ at each time point to the OD_600_ at time zero.

All assays described above were performed at least four times on at least two independent spore preparations. Data shown are averages from three replicates performed on a single spore preparation that is representative of data obtained from independent spore preparations.

### Western blot analysis

Samples for immunoblotting were prepared as previously described [72]. Briefly, sporulating cell pellets were resuspended in 100 μL of PBS, and 50 μL samples were removed and freeze-thawed for three cycles. The samples were resuspended in 100 μL EBB buffer (8 M urea, 2 M thiourea, 4% (w/v) SDS, 2% (v/v) β-mercaptoethanol) and boiled for 20 min, pelleted, and resuspended again. A small amount of sample buffer was added to stain samples with bromophenol blue. *C. difficile* spores (~1 x 10^7^) were resuspended in EBB buffer and processed as above. The samples were resolved by 7.5% (for sporulating cell analyses of CspBA and CspC) or 12% SDS-PAGE gels and transferred to Millipore Immobilon-FL PVDF membrane. The membranes were blocked in Odyssey Blocking Buffer with 0.1% (v/v) Tween 20 and probed with rabbit polyclonal anti-CspB [32], anti-CspA (a generous gift from Joe Sorg, Texax A&M University), or anti-CotA [35] antibodies and/or mouse monoclonal anti-pentaHis (ThermoScientific), anti-SleC [32], anti-CspC [30], or anti-SpoIVA antibodies [73]. The anti-CspB and anti-CspC antibodies were used at 1:2500 dilutions, the anti-SleC antibody was used at a 1:5000 dilution, and the anti-pentaHis, anti-SpoIVA, anti-CotA, and anti-CspA antibodies were used at a 1:1000 dilution. IRDye 680CW and 800CW infrared dye-conjugated secondary antibodies were used at 1:20,000 dilutions. The Odyssey LiCor CLx was used to detect secondary antibody infrared fluorescence emissions.

## Acknowledgments

We would like to thank J. Sorg for generously sharing a codon-optimized version of *cspC* and the anti-CspA antibody; N. Minton (U. Nottingham) for providing us with access to the 630Δ*erm*Δ*pyrE* strain and pMTL-YN1C and pMTL-YN3 plasmids for allele-coupled exchange (ACE); and Marcin Dembek for directly providing these materials to us and sharing his specific protocols on ACE with us. Crystallographic data collection at GM/CA@APS has been funded in whole or in part with Federal funds from the National Cancer Institute (ACB-12002) and the National Institute of General Medical Sciences (AGM-12006). This research used resources of the Advanced Photon Source, a U.S. Department of Energy (DOE) Office of Science User Facility operated for the DOE Office of Science by Argonne National Laboratory under Contract No. DE-AC02-06CH11357. The Eiger 16M detector was funded by an NIH–Office of Research Infrastructure Programs, High-End Instrumentation Grant (1S10OD012289-01A1).

